# Exploring Replay

**DOI:** 10.1101/2023.01.27.525847

**Authors:** Georgy Antonov, Peter Dayan

## Abstract

Exploration is vital for animals and artificial agents who face uncertainty about their environments due to initial ignorance or subsequent changes. Their choices need to balance exploitation of the knowledge already acquired, with exploration to resolve uncertainty [1, 2]. However, the exact algorithmic structure of exploratory choices in the brain still remains largely elusive. A venerable idea in reinforcement learning is that agents can plan appropriate exploratory choices offline, during the equivalent of quiet wakefulness or sleep. Although offline processing in humans and other animals, in the form of hippocampal replay and preplay, has recently been the subject of highly successful modelling [3–5], existing methods only apply to known environments. Thus, they cannot predict exploratory replay choices during learning and/or behaviour in dynamic environments. Here, we extend the theory of Mattar & Daw [3] to examine the potential role of replay in approximately optimal exploration, deriving testable predictions for the patterns of exploratory replay choices in a paradigmatic spatial navigation task. Our modelling provides a normative interpretation of the available experimental data suggestive of exploratory replay. Furthermore, we highlight the importance of sequence replay, and license a range of new experimental paradigms that should further our understanding of offline processing.

---

Subjects use direct experience to learn two structurally different quantities relevant to their choices: model-free values that quantify the long-run summed reward expected from performing an action; and a model or cognitive map of the environment or task they face [6, 7]. Model-free values offer a simple way of specifying a behavioural policy. Following Sutton [8], Mattar & Daw [3] suggested that replay during offline behavioural states could be interpreted as subjects employing the model to simulate potential experiences and using them to make the model-free values more accurate.

Each replay update can potentially improve a subject’s policy. Mattar & Daw [3] showed that the maximal expected improvement is achieved when the choice of state and action to replay is determined by two factors: Gain and Need (see Supplementary information). Gain quantifies the extra reward the subject expects to receive from the newly updated policy at the update state. Need is a global measure of the relevance of the update state (the strength of the successor representation there [9]) under the old policy. The combination of these factors allows replay to propagate information about reward efficiently through the environment.

However, the forms of Gain and Need in Mattar & Daw [3] assume that the model of the environment is known. Subjects are instead typically at least partially ignorant, because of incomplete initial information, forgetting or change. Exploration is thus required – and was indeed the original rationale of Sutton [8]’s DYNA architecture. Absent exploration, replay choices would be purely exploitative, and thus incomplete (Fig 1). Here, we study how replay can help generate behavioural policies which trade exploration off against exploitation in an approximately optimal way.

**Figure 1:**
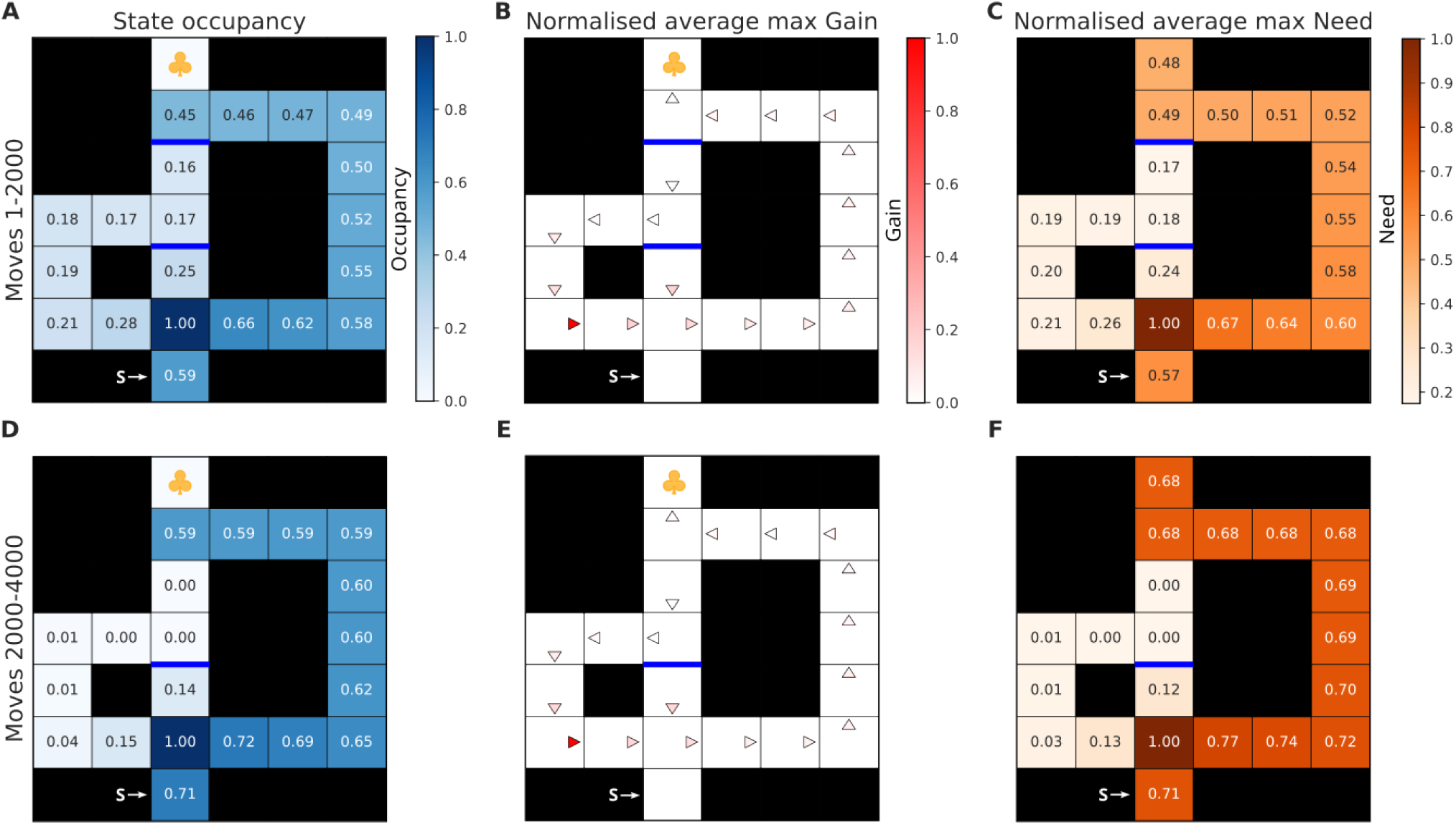
Exploitative replay can result in suboptimal behaviour. A) Normalised state occupancy of the subject during first 2000 moves of exploration and learning in the environment. The start state is located at the bottom (shown with the white letter ‘S’) and the goal state is shown with the yellow clover. The barriers are shown as opaque blue lines. Importantly, all barriers were not bidirectional, and hence could only be learnt about when attempted from an adjacent state from below. All states were visited by the subject, including those besides the barriers (darker blue corresponds to higher occupancy). B) Normalised maximal Gain that the subject estimated for the replay of each action (depicted with triangles), averaged across all 2000 moves. Only those actions for which the Gain was estimated to be positive are shown (darker red corresponds to higher Gain). The actions which the subject would replay yielded a more exploitative policy which helped the subject acquire reward at a higher rate. C) Normalised maximal Need for each state that the subject estimated, also averaged over those same 2000 moves. All values were additionally averaged over 10 simulations. Darker orange corresponds to higher Need. D-F) Same as (A-C) but for additional 2000 moves during which the top barrier was removed. Note that the estimated Gain did not change. Moreover, the state occupancy profile in D), as well as the estimated Need in F) highlight how the subject’s behaviour reduced to pure exploitation. Because of the environmental change, however, this behaviour was rendered suboptimal due to the existence of a shorter path that the subject did not discover.

There are two coarse flavours of exploration: undirected and directed [10], along with many heuristic and approximate versions of the latter. Undirected exploration comes from introducing stochasticity into choice. Although sometimes effective [11], it is typically suboptimal. Rather, exploration should be directed to reducing the uncertainty about which actions in the environment are ultimately best [12]. One standard heuristic [8] (see also [13]) is to add a form of notional exploration bonus to the outcome of actions whose consequences are uncertain.

Optimal exploration amounts to performing optimal control in a belief-state decision problem in which the physical state of the subject in the environment is augmented by the subject’s probabilistic beliefs about the environment (in our later spatial case, how likely it thinks barriers are to have been removed). This generates policies which account carefully for the longer term consequences of the resolution of the uncertainty from exploration [14, 15], trading the potential costs and benefits of doing this off against exploitation of current knowledge. Such principled accounting is radically computationally intractable, for instance because the space of possible beliefs is continuous, implying that the optimal policy can be very complex. We show how exploratory forms of Gain and Need (which extend the original notions to the belief-state decision problem) can generalise the use of replay to realise a limited version of this accounting offline.

We demonstrate the implications of our theory in a rich spatial environment specifically designed to illuminate all facets of the hard exploration problem faced by animals (and, to highlight the generality of our results, we report in the supplement the simpler case of multi-arm bandit problems). Our maze, inspired by Tolman [16], comprises three corridors which merge onto the common stem leading to the goal location (Fig 1). Those corridors differ in length, and thus an optimal reward-maximising agent (and rats [16]) would prefer the shortest corridor. However, either just the shortest, or all but the longest, path might possibly be blocked by barriers. The accompanying uncertainty provides the motivation for exploration.

For exploratory Gain: suppose that the subject is at a physical location just next to a barrier that it is uncertain is there, and is contemplating the action that might cross over and get closer to the goal (Fig S8). If the barrier is actually present, the action will fail, leading to the certain belief that the barrier is there, and no Gain. If the barrier is actually absent, this action might succeed, leaving the subject in a new location, and with a new belief that the barrier is absent. This imagined outcome is associated with high Gain, because of the implied shortcut estimated, in our account, based on the high model-free values for the new location. The net exploratory Gain comes from averaging these quantities according to the subject’s initial uncertainty about the existence of the barrier.

Exploratory Need quantifies the expected future occupancy at any given state but accounting for how the subject’s prior belief state might evolve and what it can learn in the future. However, just as the original Need, it suffers from a chicken-and-egg problem, in that if the subject adopts the purely exploitative policy of the known-to-be-open longest path, then the Need for the potential shortcut transition is zero (as the state next to the barrier is not visited). This is particularly problematic for off-policy exploration which requires visitation of states currently estimated to be unworthy. For simplicity, we make the approximation of including stochasticity in the subject’s behavioural policy (for instance, in the form of undirected exploration) such that Need is strictly positive for all possible belief states. This is achieved through applying a softmax behavioural policy [11].

The calculations of exploratory Gain and Need differ crucially from Mattar & Daw [3] in terms of generalisation. Individual physical locations (such as those next to barriers) can be visited with different beliefs about the environment. Importantly, discovering that a barrier is present/absent is information for all belief states associated with that barrier. This requires the subject to generalise the benefit of potential discoveries across multiple belief states (Fig S7).

As mentioned earlier, optimally accounting for the evolution of the subject’s beliefs is woefully intractable. We therefore incorporated an approximation [17] for the estimation of exploratory Need (see Supplementary information). The subject optimally tracks how its belief will evolve up to a limited planning horizon beyond which the residual uncertainty remains fixed. This means that the subject still maintains its subjective uncertainty about the possible futures (unlike other potential approximations [18]); however, it assumes that no new knowledge can be acquired or environmental changes take place beyond its planning horizon.

We simulated behaviour in the Tolman maze and examined the replay patterns produced as a result of uncertainty about the presence of the upper barrier (Fig 2). Note that the subject has to choose which arm to pursue at a decision point remote from the potential barrier location. There is thus substantial cost for exploration: the subject has to have sufficient belief that the barrier is open – otherwise the potential benefit of exploration (i.e., discovering a shortcut) would not exceed the cost of deviating from the current behavioural policy (i.e., its current reward rate) [19].

**Figure 2:**
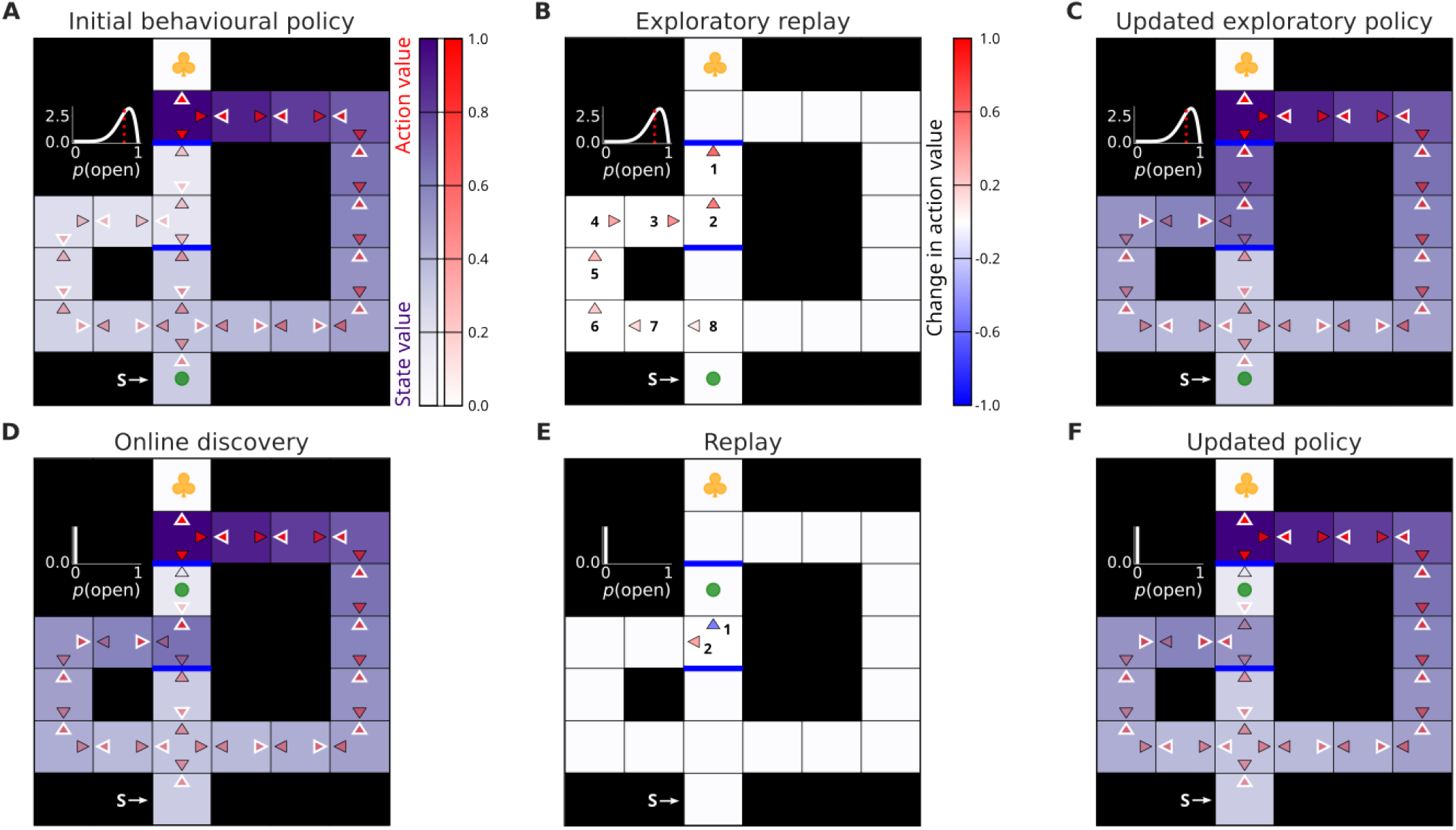
Exploratory replay leads to online discoveries, but potentially inadequate promulgation. A) Prior state of knowledge of the subject. The intensity of the (red-scale) colour of each action arrow shows the respective model-free *Q*-values. Collectively, the action values represent the subject’s model-free behavioural policy (i.e., the subject is more likely to choose actions with higher estimated *Q*-values – which at each state are highlighted with white outlines). Similarly, the states are coloured according to the maximal model-free *Q*-value at each state (which corresponds to state values, shown in purple). The inset next to the top barrier indicates the subject’s prior belief about its presence (for the other barrier, the subject was certain that the path was blocked). The red dotted line in the inset shows the expected probability that the barrier is absent. The subject itself (green dot) is located at the start state. The goal state with reward is denoted with the yellow clover. B) Changes in the subject’s model-free policy occasioned by exploratory replay updates. The numbers next to each action arrow indicate the order in which the replay updates were executed. C) New model-free policy which resulted from exploratory replay updates in B). Note how the action values now indicate that the subject should go towards the upper barrier (highlighted with white outlines). D) After pursuing the exploratory policy, the subject attempted to cross the top barrier; unfortunately, the barrier was found to be present – this is indicated by both the subject’s model-free *Q*-value associated with that action which was learnt online, as well as its new belief. E-F) Same as in B-C) but after the online discovery of the present barrier in D). The first replay choice of the subject correctly propagated the negative value of the present barrier to the immediately preceding state. However, as opposed to propagating this information deeper towards the start state, and hence correcting the exploratory policy in the light of the new information, the next replay choice of the subject made it more likely to visit an adjacent state which still contained the previously propagated exploration bonus, and hence had a high value that was erroneous given the subject’s new knowledge.

Here, the subject’s uncertainty resulted in consecutive replay updates which originated at the potential barrier location and progressed towards the subject’s location in a reverse manner (Fig 2A-C). Those replays propagated the value of exploring the barrier towards the subject’s current location, and the resulting new model-free behavioural policy indicated exploration was worthwhile (Fig 2C). As just discussed, the extent to which the subject was uncertain determined how large was the exploratory bonus that reached the subject’s current state – and thus produced policies with different incentives for exploration (Figs S5 and S6).

Resolving uncertainty can often result in unfortunate outcomes, for instance if the barrier is found actually to be present (Fig 2D). If this happens, it is important for the subject to correct the full exploratory policy that had led to the discovery in the light of the negative information it acquired. We find that in our simulated Tolman maze, single-action replay updates do not handle this appropriately: the discovered value of the present barrier does not propagate deeply enough towards those states which had been updated with the exploratory bonus of the obsolete belief (Fig 2E-F). This is because single-action updates are myopic: the estimated benefit of a single-action update does not account for how that update can affect the benefit of potential future updates. This problem does not arise if the shortcut is found to be available, or in stationary environments with monotonic value structures, since then the replay naturally spreads the (correct) good news in backwards sequence [20].

One plausible solution is to consider the benefit of simultaneously updating a sequence of actions, as opposed to relying solely on updates at single states. This benefit combines Gain, that accumulates with the propagated policy changes (provided that all those changes result in policy improvements), as well as Need along that sequence of actions. We found that sequence replay results in deep propagation of the value of a discovered barrier, along the whole chain of actions which had previously been endowed with the exploration bonus (Fig 3).

**Figure 3:**
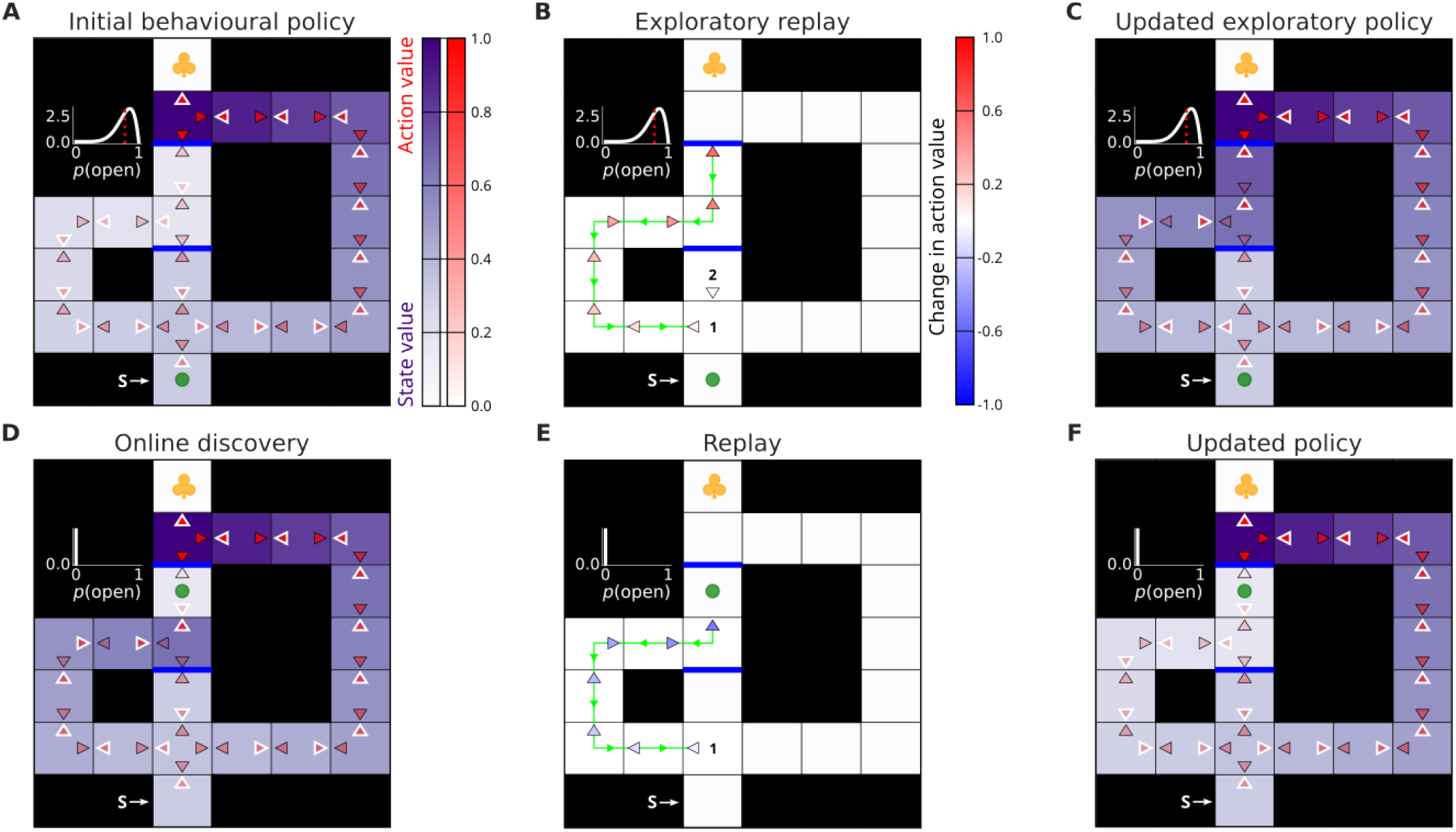
Sequence replay helps deep value propagation. The layout of the figure is the same as in Fig 2. A-C) Show the subject’s initial and uncertain state of knowledge, changes to the online behavioural policy occasioned by exploratory replay, and the new updated exploratory policy due to such replay, respectively. The crucial difference being that the replay in B) was a sequence event – i.e., the whole chain of actions was updated simultaneously (the actions which were updated in the replayed sequence are linked by a green line; the green triangles along that line additionally indicate the reverse direction of the replayed sequence). D-F) Again, the subject discovered the top barrier, learnt about its presence online and engaged in replay to recompile its model-free behavioural policy in the light of the negative information. Note how, in this case, sequence replay in E) resulted in deep propagation of the value of such information all the way towards the start state. The sequence replay thus enabled the subject to correct its exploratory policy appropriately as shown in F).

There is one further aspect of the data on exploratory replay: experimental evidence implicates the hippocampus in constructing replay sequences through previously unexplored spaces [21, 22]. In our account, this corresponds to replay in potential future belief states which the subject has not visited yet but imagines encountering. We manipulated the barrier configuration in our maze to produce a corridor segment in the central arm with both sides occluded by barriers (Fig 4A-C). Examining the replay patterns chosen by the subject due to uncertainty about the presence of both barriers revealed sequence replay in the corridor. Such replay propagated the exploratory value of learning about the possibility of entering the corridor (resolving uncertainty about the bottom barrier; Fig 4B bottom), exiting it (learning about the top barrier; Fig 4B top) and ending up in a state close to the goal. Similarly, we simulated the experiment from Ólafsdóttir *et al*. [22] which resulted in the ‘preplay’ of the goal-cued (but not uncued) arm prior to experience (Fig 4D).

**Figure 4:**
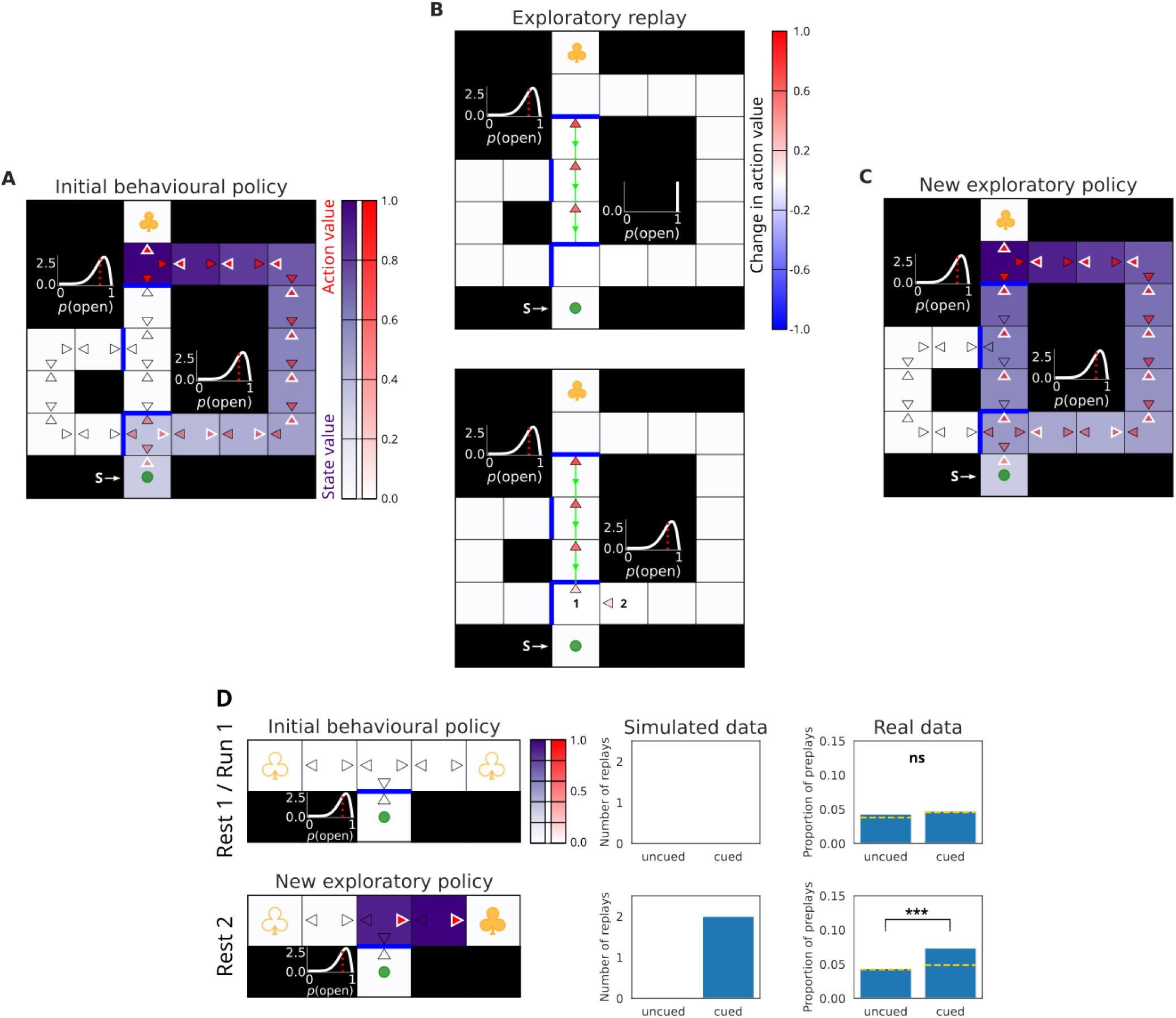
Replay in a blocked corridor. A) Initial state of knowledge of the subject. Note that the model-free *Q*-values in the blocked corridor are all initialised to 0, thus mimicking the subject’s inexperience with the segment. The subject’s belief state comprised its uncertainty about the presence of the top and bottom barriers that create the corridor. B) Replay choices of the subject due to its initial and uncertain state of knowledge. Note that the sequence replay event was performed across two different belief states: action updates inside the corridor (top) corresponded to a different belief state since they followed the potential transition through the bottom barrier which the subject had to first learn about (bottom). C) New exploratory policy occasioned by the replay updates in B). D) Same setup as above, but simulating offline rest replay in the T-maze experiment from Ólafsdóttir *et al*. [22]. The top row shows the initial state of knowledge of the subject. In the actual experiment, ‘Rest 1’ replay events were measured before the animals’ experience of the environment, and during ‘Run 1’ they explored the central stem which was blocked by a see-through barrier. In ‘Run 1’, none of the arms contained a visible reward (which are depicted with unfilled yellow clovers). No detectable replay was observed in the two arms during the ‘Rest 1’ condition. ‘Rest 2’ replay events were measured during a rest period after a visible reward was placed in the ‘cued’ arm (filled yellow clover) but before the animals could experience it (i.e., before the barrier was removed). Note that we rendered the see-through barrier as potentially permeable (as reflected in the subject’s uncertain belief) due to which the subject could contemplate during rest the possibility of crossing it and obtaining the reward. The bottom row shows the resulting exploratory policy after the subject was allowed to replay with the knowledge of the reward in the cued arm. This new policy resulted from replay in the cued (and not uncued) arm. Note that, as in B), such replay was performed in a different belief state (corresponding to learning that the barrier was open) than the subject’s prior belief state, and thus could potentially only be detected after the actual experience. Data from Ólafsdóttir *et al*. [22]. Yellow dotted lines show chance detection level. ns, not significant; ***, *p <* 0.001.

Some of the most important facets of learning in the brain involve building inverse models: this characterises bottom-up, recognition, models of sensory processing in cortex [23]; the maintenance and expansion of the relationship between cortical and hippocampal representations in memory [24– 26]; and the determination of policies that maximise reward and minimise punishment given information about the environment [8]. Offline processing, evident in replay, offers a way of building and refining inverse models of all these forms without disturbing ongoing behaviour. However, to determine good policies, it is not enough to build an inverse model based on just current information; active observers have the obligation to collect new information too, and balance this against exploitation. This obligation can be satisfied by inverting a more sophisticated model of the environment that includes uncertainty; here, we showed how to conceive of (reverse) replay as performing this inverse. This provided new insights into the nature and structure of offline activity – for instance surfacing the importance of sequence replay, as well as predictions for new experimental paradigms.

## 1 Methods

We treated the (navigational) decision-making problem in our variant of the Tolman maze as a partially observable Markov Decision Process (POMDP). The subject was designed in the spirit of the DYNA architecture, such that online decisions were made according to the behavioural model-free policy, and offline planning was used for additional training of the model-free controller. The subject was endowed with a probabilistic belief about the existence of barriers in certain locations in the maze; every decision (real or imagined) therefore transitioned the subject to a new belief state which comprised the subject’s physical state, as well as its updated posterior belief, which became its new prior belief. For planning (replay), the subject considered how its belief state would evolve up to a fixed horizon. The value of each imagined belief state was approximated with the subject’s model-free *Q*-values at the corresponding physical location. Moreover, we considered just three possible beliefs for the existence of each barrier: the initial uncertainty (which can be continuous), and either certain presence or absence. The priority of each replay update was determined by the expected long-run improvement to the subject’s current belief state engendered by each potential replay update. The replay updates were executed until the expected improvement was estimated to be below a fixed threshold. For sequence replay updates, the maximal length of each potential sequence was limited to the distance from the start state to the uncertain barrier. We report a more detailed theoretical account of our modelling in the Supplementary information.

## 2 Supplementary information

### Theory background

#### Reinforcement learning

In reinforcement learning (RL) [6], subjects learn to make appropriate decisions in order to maximise expected gains and minimise potential losses. Learning proceeds through interaction with an environment which supplies a sparse learning signal. The environment is typically formalised as a Markov Decision Process (MDP), which is a tuple ⟨ 𝒮, *𝒜, 𝒫, ℛ, γ*⟩ where S is the set of states, A is the set of actions available at each state, 𝒫 : 𝒮 × 𝒜 × 𝒮 → [0, 1] is the Markov transition kernel which specifies the transition probabilities between states given an action, ℛ : 𝒮 → ℝ is a bounded reward function which comprises the learning signal, and *γ* ∈ [0, 1) is the discount factor which determines the appetitiveness of delayed rewards.

The subject’s behaviour in an environment is governed by its policy, *π* : 𝒮 × 𝒜 → [0, 1], which, for every state, outputs a probability distribution over the set of available actions. At each time step, the subject interacts with its environment and receives the reward signal. The (possibly infinite) discounted collection of rewards the subject accrues along a trajectory of decisions is called the return. One main goal for a reinforcement learning subject is to predict the expected rewarding consequences of following policy *π* starting at a state *s*. This can be written as

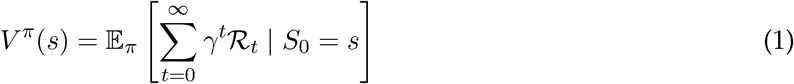

A closely related task is instead to estimate the expected return for performing some action *a* in a given state *s*, in which case they are referred to as *Q*-functions:

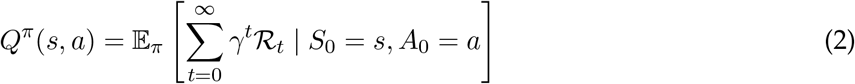

The second main goal is to learn an optimal policy, *π*^*^, which for any starting state *s* prescribes how to maximise the expected return:

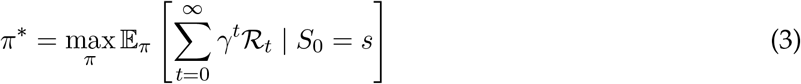

An MDP need not have a unique optimal policy. However, the optimal value function 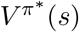 and 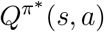 functions are unique. In particular, any action 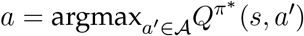 can be chosen.

#### Model-free control

Several algorithmic approaches exist to solving the problem of optimal control in RL tasks. One popular example is *Q*-learning [27], which is an important and widely used algorithm for learning the optimal 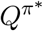 -function. It belongs to a more general class of model-free temporal difference algorithms which, after every experienced interaction with the environment, successively update their value function estimates based on the encountered reward prediction errors. Specifically for *Q*-learning, the update rule at iteration *n* is:

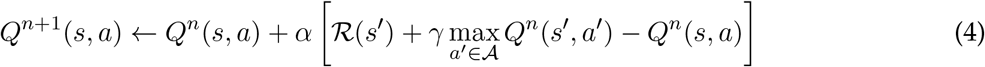

Here, the *Q*-value estimate is updated towards the difference (or prediction error) between the initial estimate, *Q*^*n*^(*s, a*), and the sum of the observed reward at the next state reached and the discounted maximal *Q*^*n*^-value at that state, ℛ (*s*^′^) + *γ* max_*a*_′_∈𝒜_ *Q*^*n*^(*s*^′^, *a*^′^), weighted by the learning rate, *α*. Note that the action that optimises *Q*^*n*+1^(*s, a*^′^) at *s* might be different from the one used in equation 4 that optimised *Q*^*n*^(*s, a*^′^)

The *Q*^*n*+1^-values themselves can be used to determine a policy, for instance:

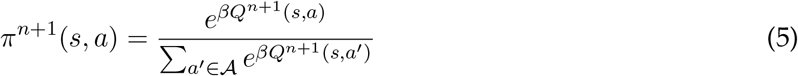

where *β >* 0 is an inverse temperature parameter that controls how deterministic is *π*^*n*+1^. Since *π*^*n*+1^(*s, a*) favours actions with higher *Q*^*n*+1^-values, it tends to be better than *π*^*n*^(*s, a*) in terms of expected return. The remaining stochasticity is a crude method for arranging a mix of exploration and exploitation.

#### Model-based control

A different solution is to learn a model of the environment which can then be used to perform prospective *planning* of the actions to execute. Value functions can also be acquired using the recurrent Bellman equation [28], for instance:

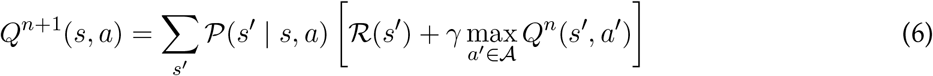

Here, the recurrent relationship between the successive states allows the subject to make use of its knowledge of the transition structure of the environment (the model 𝒫) to propagate the information about future rewards towards its current situation or state in the environment. If the subject does indeed know the model (also including ℛ), then various forms of planning can be used to compute the long-run consequences associated with the available actions at decision time and make a far-sighted and informed decision. Value iteration [28] is one example planning algorithm which iteratively performs synchronous updates (for all states and actions in each sweep) specified by Equation 6. Such updates are also called Bellman backups because of the application of the Bellman equation. Given a perfect model of the environment, 𝒫, such procedure is guaranteed eventually to converge to the optimal value function.

#### DYNA and prioritized sweeping

There is evidence for the use in animals, and the utility in artificial agents, of both model-free and model-based control [29]. This poses obvious questions about their arbitration and integration [5, 7, 13]. One important suggestion for integration is that information could be transferred from the model that the model-based controller possesses into the model-free controller, so that the latter can provide better informed choices.

In RL, the most common version of this process is known as experience replay [30], and lies at the heart of many successful algorithms [31]. Although, as we will discuss later, it was originally designed for the purpose of exploration, the so-called DYNA algorithm [8] has been used to underpin this process. In DYNA, an agent learns model-free value functions online by direct experience with the environment, as well as learning the model of that environment. During offline states, DYNA uses its learnt model to sample possible transitions and rewards, which are then used to perform further training of the model-free value functions to perform a more effective form of model inversion.

Given this overall structure, it becomes natural to consider which transitions or rewards should be sampled from the model (or replayed). One important algorithmic notion is prioritized sweeping [20], in which replays are chosen in an order that effects a form of optimal improvement in the model-free value functions.

#### Gain and Need

Mattar & Daw [3] synthesised the ideas of DYNA and prioritised sweeping and proposed a principled, normative scheme for the ordering of planning computations. They suggested that each replay experience corresponds to a Bellman backup (Equation 6) which uses information from a generative model of the environment to update a specific model-free state-action value.

Mattar & Daw [3] observed that what is important about an update at a state (which could be distal from the current state of the agent) is whether it changes the subject’s behavioural policy. For example, performing a planning computation at state *s*_*k*_ corresponds to changing the model-free value for action *a*_*k*_ at that state. Such a change is significant if the agent’s behavioural policy changes at *s*_*k*_; the agent can then estimate the consequence of that change for the expected return from its current state or a start state.

Mattar & Daw [3] showed that the subject can calculate how a replay update to action *a*_*k*_ at state *s*_*k*_ changes the amount of reward it can obtain in the future starting from a potentially different state *s*. By decomposing the difference in the subject’s model-free value function estimate before and after the policy update occasioned by such replay update, 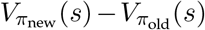, Mattar & Daw [3] showed that this expression can be written as:

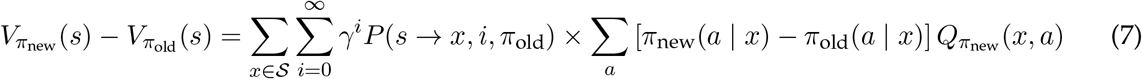

Furthermore, by assuming that each individual replay update to the model-free value of action *a*_*k*_ results in a policy change at a single update location, *s*_*k*_, equation 7 can be simplified into the product of Gain and Need, which Mattar & Daw [3] termed the expected value of a backup 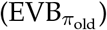:

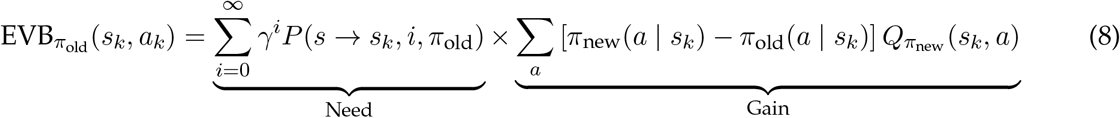

Gain quantifies the expected local improvement in the subject’s behavioural policy at state *s*_*k*_ as a result of the replay update. Thus, Gain is higher for those replay updates which result in greater policy changes at the update state. Need, on the other hand, quantifies how likely is the subject to visit the update state in the long run, given its model of the environmental transition dynamics and behavioural policy before the update.

In rodents, the hippocampus is a structure known to be involved in aspects of model-based control [32–34]. Mattar & Daw [3] suggested that the reactivation of sequences of behaviourally-relevant experiences during quiet wakefulness and sleep for which the hippocampus is well known [35] is an expression of this sort of prioritized replay. They thereby explained a wealth of experimental findings on the selection of replay experiences in rodents [32, 33] as well as humans [5, 36].

#### Exploration

As discussed in the main text, exploration in MDPs can be accomplished by the use of heuristics which estimate the amount of the subject’s (in)experience with its environment. One such celebrated heuristic is based on the ‘optimism in the face of uncertainty’ (OFU) principle which posits that actions whose outcomes are uncertain should receive a sort of exploration bonus which would encourage the subject to pursue them. Sutton [8]’s exploration bonus indeed took that form:

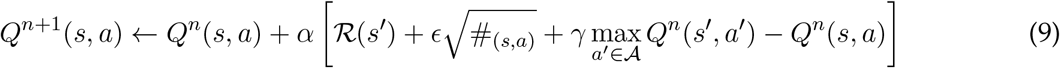

Improved exploration in DYNA (also known as DYNA-*Q*+) was achieved by updating its model-free *Q*-values according to Equation 9 during offline planning. Here, #_(*s,a*)_ is a count-based heuristic which grows with the number of time steps since that state-action pair had last been attempted, and *ϵ* is a free parameter which controls the amount of influence this uncertainty bonus has on the *Q*-value update. By using this update rule, actions which have not been tried for an extended period of time come to look more appealing, which happens to be particularly useful in dynamic environments with unsignalled changes.

Note that by virtue of the *Q*-learning update rule (Equation 4), the exploration bonus awarded to a distal state-action pair (Equation 9) propagates towards state-actions which lead to it, hence encouraging off-policy exploration. The bonus itself, however, is myopic, since it does not reflect the benefit of learning about the uncertain state-action in the first place.

Optimal exploration, on the other hand, entails a more careful evaluation of how resolving one’s uncertainty may be useful in the long-run and whether the acquired knowledge would be of any use for subsequent exploitation. Such thorough evaluation requires the subject to maintain an explicit model of its uncertainty and what possibilities abound.

#### Partial observability

The classical MDP formalism assumes that the subject knows the model of the environment with which it interacts. It does not, however, capture the ignorance that subjects (at least partially) face when learning about their environments. Such ignorance can be treated as a form of incomplete information which the subject can (at least to some extent) complete with experience.

Partially observable Markov Decision Processes (POMDPs) are a generalisation of MDPs in which the subject can lack direct access to some knowledge that is required to learn a good policy. For instance, the subject can be ignorant about the state it occupies because instead of perfect information from the environment it receives noisy and ambiguous observations; equally, the subject can be uncertain about the transition dynamics that govern its movement through the environment.

Each observation in a POMDP therefore grants the subject a piece of information which it can use to update its knowledge about the environment in an optimal manner. A sequence of observations the subject collects is formally referred to as *history*. Critically, the subject’s policy in a POMDP depends on its full history of observations, since this history determines its state of knowledge about the environment, and thereby determines the decisions it ought to make. The dependence on history violates the Markovian assumption (which requires that future transitions and rewards are statistically independent of the history, given the present state), and POMDPs are therefore not amenable to classical MDP solutions.

Instead of keeping track of all encountered observations the subject can maintain a sufficient statistic of the entire history. This sufficient statistic is called the subject’s *belief*, and it concisely summarises the knowledge that the subject has acquired. With each new observation the subject can optimally update its beliefs in the light of new information. Beliefs can be viewed as a new, subjective, state for a decision problem; they do satisfy the Markov property, and so it is possible to formulate POMDPs as MDPs where each state of the process is the subject’s belief.

A belief MDP is therefore formally defined as a tuple ⟨ ℬ, *𝒜, 𝒯, ℛ, γ*⟩ where B is the (continuous) set of belief states, 𝒜 is the set of actions, 𝒯 : ℬ × 𝒜 × ℬ → [0, 1] is the (Markov) belief transition kernel, ℛ : ℬ → ℝ is a bounded reward function, and *γ* ∈ [0, 1) is the discount factor. Thus, as opposed to the original MDP formulation, in belief MDPs the subject transitions through augmented belief states. For our matters, each belief state, *b* = {*s* ∈ 𝒮, *P* (𝒫)}, encompasses the subject’s physical location in the environment, *s*, as well as its probabilistic model of uncertainty, *P* (𝒫), about the presence/absence of barriers at several locations.

The formalism of belief MDPs permits the construction of policies which optimally trade-off exploration and exploitation [37]. To see this, consider the case that the subject is uncertain about the state transition model 𝒫, and therefore maintains a prior belief *P* (𝒫). Firstly, the probabilistic belief allows the subject to learn optimally upon receiving observations from the environment – in the case of transition uncertainty, by noting which state each transition leads to. This is accomplished by calculating a posterior belief using Bayes’ rule. For instance, after observing a transition from state *s* to *s*^′^, an optimal belief update corresponds to:

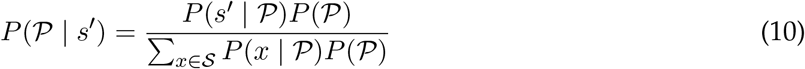

Note that a general POMDP formalism typically involves an observation function whereby the subject has no direct access to the state of the world, and it therefore receives noisy observations which lead to uncertain state estimates. In our setting, the subject has direct access to its physical state in world; however, the transition structure is non-trivial in the sense that it can change without the subject being aware of such changes taking place. The subject’s uncertainty can result from either the subject having an explicit probabilistic belief of how the transition dynamics might change in the course of a task, or, alternatively, because of forgetting, which can be thought of as a heuristic version of the former.

Secondly, the subject can plan the future possibilities by making use of its uncertainty and allocating the prior probabilities to each of the considered outcomes. Those outcomes, in turn, result in more potential learning which the subject also accounts for by performing the same updates as in Equation 10 but for simulated futures (those transitions are governed by the belief MDP transition function, 𝒯). This allows the subject to foresee the long-run consequences associated with each exploratory decision and whether it can potentially result in better future return.

### Model description

#### Replay updates

The subject makes use of its transition model as well as the associated uncertainty to envision the possible evolution of its belief. This can be visualised as a planning tree which is rooted at the subject’s current belief state, *b*_*ρ*_. The subject considers all possible actions from this root node, and adds additional nodes for each new belief state that results from applying those actions (according to the belief transition model, 𝒯) – this corresponds to adding a single step horizon to the planning tree. Applying the same procedure to all nodes at the new horizon further deepens the tree and expands the planning horizon.

Similarly to physical states in MDP problems, each belief state can have an associated value which reflects how much reward the subject expects to obtain by being in that belief state and acting according to some policy. Those values, however, are initially unknown to the subject, and the reason for performing replay updates in the belief tree is to propagate the value information from future belief states to the subject’s current belief state. Since belief states are continuous, we restrict the subject’s planning horizon to a fixed depth. This means that belief states containing reward may be beyond the subject’s reach. However, the subject’s model-free system is likely to have an estimate of how valuable each physical location is. Therefore, the model-based value of each action *a* at every belief state *b* = {*s, P* (P)} in the planning tree, which we refer to as 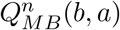, is initialised to the subject’s model-free estimate of the value of performing this action at the physical location in that belief state, 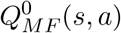.

When performing replay updates, the subject considers the effect of each action at every belief state in the tree rooted at its current belief state. For example, when considering the effect of action *a* at belief state *b* = {*s, P* (𝒫)} which attempts to cross a potential barrier, the subject accounts for the possibility of transitioning into one of two new belief states: 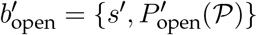, which corresponds to the fortunate outcome of discovering that the barrier is absent, and 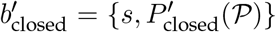, which corresponds to the unlucky outcome of the barrier being present. The value associated with executing action *a* at belief state *b* is updated towards the estimated values of the next belief states:

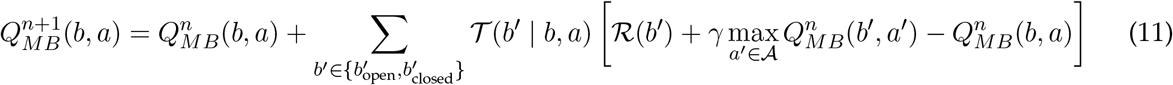

Here, the belief transition model, 𝒯, describes how the subject jointly transitions through physical states and its beliefs about the barrier configuration. Moreover, for brevity, we will refer to the set of belief states that the subject can reach by applying a single action at a belief state as the children set of that belief state, denoted as *C*(*b, a*) ∈ ℬ. For the example above:

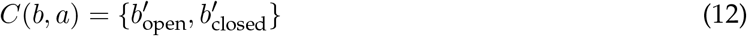

#### Gain and Need in belief space

We consider optimising the prioritisation of replay updates (Equation 11) in the subject’s belief space. We follow the suggestion of Mattar & Daw [3], whereby the priority of each update is determined by the expected improvement to the subject’s behaviour at its current belief state. By applying the same value decomposition as in Mattar & Daw [3], we define 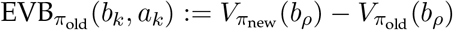, where 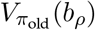 is the value the subject estimates for its current belief state, *b*_*ρ*_, under the old behavioural policy before the potential update, and 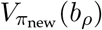 is the estimated value of the subject’s current belief state under the new policy implied by the potential update. The effect of policy change engendered by a replay update to action *a*_*k*_ at some (potentially distal) belief state *b*_*k*_ can be expressed as:

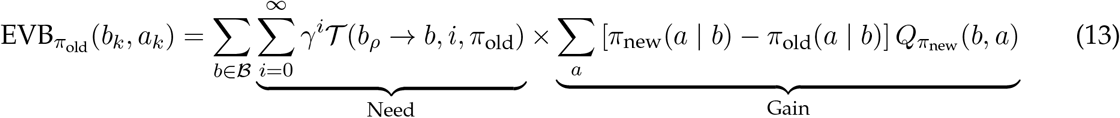

Importantly, we do not assume that the effects of replay updates are localised to individual states (as in Equation 8), which allows the subject to account for broad generalisation across multiple belief states (see below) when calculating the expected benefit of each replay update. The Gain term associated with a replay update quantifies the expected local improvement in the subject’s behavioural policy at the update belief state engendered by that replay (Equation 11). Gain therefore favours those replay updates which result in large improvements to the subject’s model-free decision policy.

Need, similarly to Mattar & Daw [3], quantifies the frequency with which the subject expects to visit the update belief state according to its old behavioural policy, *π*_old_. As discussed before, in belief MDPs, subjects engage in continual learning which means that with every visit to the same physical location the subject, in general, will have a different belief about the transition model. This allows the belief space version of Need to account for all possible future learning that can take place (however, for computational purposes, we limit the subject’s horizon – see below).

One critical consideration is that of the dependence of Need on the old behavioural policy of the subject, *π*_old_, which tends to prioritise portions of the state space the subject already expects to visit. Thus, even if the subject was informed about a distal change in the transition structure which its current policy does not prescribe to visit, Need at those locations would still be zero. It is therefore important to include stochasticity (for instance, in the form of undirected exploration) into the subject’s behavioural policy which generates Need to allow for off-policy replay choices. This motivates our choice of the softmax behavioural policy which ensures that Need is positive for all potential belief states. Note that such design is common to most planning algorithms as it ensures adequate exploration of the state (and belief) space when performing planning computations [38, 39]. Below we additionally explore how the subject’s behavioural policy affects it replay choices.

As for Mattar & Daw [3], we set a threshold on the minimal 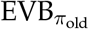 value required for an update to be executed. This threshold can be thought of as accounting for a form of opportunity cost by balancing the trade-off between planning to improve the policy and immediately acting to collect reward [19], hence helping to subject to avoid being permanently buried in thought.

#### Generalisation

The notable difference between our belief space decomposition and that of Mattar & Daw [3] is the inclusion in equation 13 of the outer sum over the space of beliefs, ℬ. This critical difference enables the subject to account for a broad generalisation across multiple belief states when considering the effect of a single action update at an individual belief state (Fig S7).

In the original formulation of Mattar & Daw [3], the accumulated benefit of policy change at a physical state arises due to the repetitive visitation of that state that the subject envisions according to its behavioural policy and its model of the environmental transition dynamics. This form of Need corresponds to an approximation based on the past experience of the subject which assumes that no further knowledge can be acquired. Our formulation allows accounting for future occupancy based on the potential future learning that can take place in the environment. Such accounting requires the subject to generalise information learnt at individual physical states across multiple potential beliefs at which the subject can re-visit that physical state in the future (Fig S7).

In general, each belief state in continual learning tasks (unless there is forgetting) can be visited at most once since after every transition the subject potentially acquires information, and therefore updates its prior belief which constitutes a different belief state (this is true especially for Bayes-adaptive MDPs; [2]). The POMDP framework can be adapted such that this need not always be the case, since for instance in the Tolman maze which we consider here, the subject maintains uncertainty about the presence of barriers at certain locations, and this uncertainty can only be reduced so long as the subject actually attempts to cross those barriers. Therefore, when the subject transitions through those states which it is perfectly certain about there is no information gained as regards its belief about the barrier configuration, and thus the physical state is the only constituent of the belief state which changes (hence the subject can in fact visit a physical location with the same belief multiple times). Although this is exactly how we modelled our subject’s uncertainty about its environment, the replay formalism we developed here is more general and applies also to settings in which beliefs change after every transition or observation.

In the presence of forgetting, the replay structure might be different since the subject would need to optimally account for those belief states which it expects to visit again. This, however, will depend of the specific form of forgetting, and the resulting belief states which the subject would have to represent in the planning tree. Our general formalism of replay prioritisation can account for this, but in the present work we do not consider it.

#### Sequence replay

Sequence replay corresponds to updating a whole sequence of consecutive actions, as opposed to performing individual greedy action updates one at a time. For example, consider two consecutive actions *a*_1_ and *a*_2_ at belief states *b*_1_ and *b*_2_, respectively. The order in which those two replay updates are executed depends on the expected value associated with the two possibilities. In the spatial domain (or other domains with clear ordering) one order would typically be interpreted as a reverse reactivation, and the other as forward. Moreover, the expected value of performing forward and reverse sequence updates will, in general, differ (see below). A sequence update to the two example actions corresponds to updating one action according to:

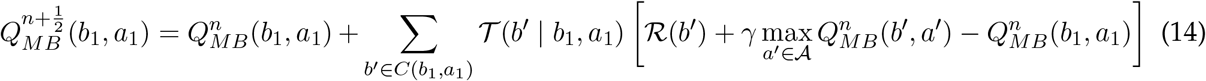

where the sum is over the set of next possible beliefs (as in equation 12). The fractional notation 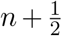 emphasises the fact that within a single iteration of replay multiple actions can simultaneously be replayed in a sequence, since in the current example with two actions there are two executed updates between iterations *n* and *n* + 1.

The second action is then updated in the same way to generate *Q*_*MB*_; however, in the case of reverse replay, *b*_1_ ∈ *C*(*b*_2_, *a*_2_), and therefore the 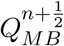-value of one of its children beliefs *b*^′^ ∈ *C*(*b*_2_, *a*_2_) will have already been updated. The size of the value update to action *a*_2_ at belief state *b*_2_ therefore depends on the update to action *a*_1_ at belief state *b*_1_. This is also reflected in how the expected value of sequence replay is calculated – which is the reason for why the benefit of sequence replay can be larger than that of single action updates. If we define ℳ _*N*_ = {(*b, a*)_*i* 1,…,*N*_} as the candidate set containing *N* belief state-action pairs to be potentially updated in a sequence replay event, then the expected benefit of that sequence replay is calculated as:

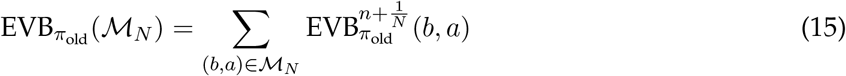

Note that, in the case of reverse replay, each individual 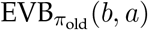 in Equation 15 quantifies the benefit of updating action *a* at belief state *b* with a value that is propagated towards it along the sequence of actions that had also been updated. This is not the case for forward replay where each action is updated only towards the expected value of its children belief states (with the exception of cyclic domains; however, as we report below, we restrict all sequences to be acyclic); however, even in the case of forward replay the benefit of replaying the whole sequence will still, in general, be higher because of the summed benefit of all updates along the entire sequence (see below).

Replayed sequences can be of arbitrary lengths. Moreover, the longer the sequence, the more the estimated expected benefit will be, in general. The natural question therefore arises concerning the termination of sequences. We do not address this issue in the current work and assume that sequences link together critical decision points – in the Tolman maze, for instance, this corresponds to the sequential replay which originates at a potential barrier location and progresses towards the intersection in front of the subject’s start state.

Another consideration is computational: the theory that Mattar & Daw [3] proposed is normative and does not prescribe how both Gain and Need can possibly be estimated in a psychologically credible way. Sequence replay is even more computationally prohibitive because of the number of potential sequences that can be replayed. In the present work, we similarly report a normative result describing which sequences (out of all possibilities up to a fixed length) should be replayed. How the brain manages to reduce the sample complexity of sequence replay thus remains an open and challenging question which we leave to future work.

#### Simplified example: Bayesian bandits

Stationary, multi-arm bandit (MAB) problems offer the simplest test bed for examining exploration in belief spaces, and we therefore provide simulation results of replay prioritisation in a class of MABs. A typical MAB problem consists of a finite set of *K* arms, 𝒜 = {*a*_1_, …, *a*_*K*_ }, which are the equivalent of actions in sequential decision-making problems. In each of the infinitely many trials, the subject is faced with a choice to pull one of the available arms. Each of the *K* arms, say *a*_*k*_, if chosen, has a certain probability, *μ*_*k*_, of paying the subject off with a binary reward (1 with probability *μ*_*k*_ and 0 with probability 1− *μ*_*k*_). One typical goal of subjects in MAB problems is to realise a sequence of arm choices so as to maximise the total discounted reward.

MAB problems are well-studied and, under certain assumptions about the reward distribution, optimal policies can be derived (such as the Gittins index [14]). Importantly, the payoff probabilities associated with each arm are initially unknown to the subject. This makes exploration in MAB problems worthwhile even if the expected return for the arm concerned is low, since if the arm is found actually to be good, then it can be consistently exploited in the future. Furthermore, MABs lack physical states, since in each trial the subject is faced with the same selection of arms irrespective of its choices in the preceding trials. The lack of physical states and the necessity of exploration makes MABs a perfect case study for our replay prioritisation, which we detail below.

We focus on a 2-arm bandit task with binary outcomes, in which on each trial, the subject has to choose between two arms, *a*_1_ and *a*_2_, which have unknown payoff probabilities, *μ*_1_ and *μ*_2_, respectively. The subject models its uncertainty about the payoff probability of each arm with a probabilistic prior belief which introduces subjective belief states, *b* = {*p*(*μ*_1_), *p*(*μ*_2_)}. Just as in the Tolman maze example considered above, a probabilistic model of uncertainty allows the subject to learn optimally about the payoff distribution of each arm after receiving feedback from the bandit in the form of a reward signal.

We model the subject’s uncertainty about each arm’s payoff probability using the Beta distribution. This particular parametric form is very convenient since the Beta distribution is a conjugate prior for the Bernoulli distribution (which is the reward distribution of each arm). The Beta distribution has two parameters, *α* and *β*, where *α* is typically interpreted as the number of success trials (received a reward of 1) and *β* as the number of failed trials (received a reward of 0). After *N* choices of arm *a*_*k*_, the Bayesian update (Equation 10) to the prior distribution parameters (i.e., the subject’s belief state) due to a new observation corresponds to:

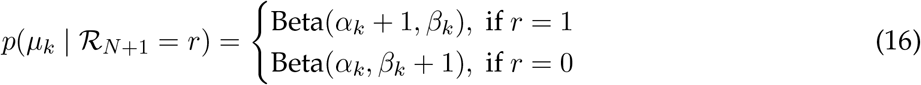

where *α*_*k*_ + *β*_*k*_ = *N*.

The subject can make use of its model of uncertainty to plan ahead how the choice of each arm will affect its belief. We visualise this as a planning tree in Fig S1A. The tree is rooted at the subject’s current belief state, *b*_*ρ*_, and each action (choosing arm *a*_1_ or *a*_2_) can transition the subject into two new possible belief states: one which corresponds to an imagined success trial and another corresponds to an imagined failure trial. Applying actions to belief states deepens the tree and expands the subject’s planning horizon. Note that the subject’s planning horizon is limited to a fixed depth (Fig S1). This is because belief states are continuous, and building the entire tree of all possibilities is intractable. In our example, the subject therefore only considering how its belief will evolve up to several steps into the future.

Analogously to the Tolman maze, each belief state has an associated value. Replay updates in the tree correspond to updating the value of each belief state towards the expected value of the beliefs of its children at one horizon deeper in the tree (Equation 11). We initialise the value of each action *a*_*k*_ in every belief state *b* in the tree to 0, except for the belief states at the final horizon whose values are initialised to the immediate expected payoff the subject expects to receive in that belief state by choosing action *a*_*k*_, which corresponds to 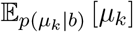.

Similarly to the belief MDP, we define the priority of each individual replay update in the subject’s belief space as the expected value of the associated backup (EVB). That is, for a potential replay update at belief state *b*_*k*_ to the value of action *a*_*k*_, the expected value of that update, 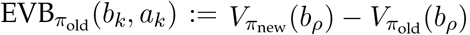, which quantifies the expected improvement to the value of the subject’s current belief state, decomposes into the product of Gain and Need:

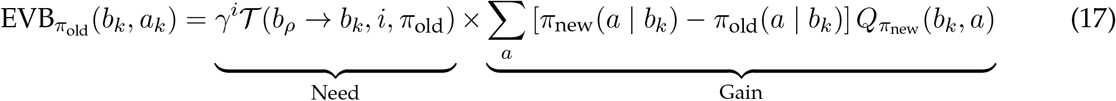

The MAB instance of Gain is very similar to that of a more general belief MDP Gain discussed earlier. The crucial difference, however, is that Need does not accumulate at any individual belief state (which is why we refer to it as non-cumulative Need). This is because there are no physical states which can be re-visited in each episode, and each belief state in an MAB can be visited at most once (provided there is no forgetting involved) due to the continual learning nature of the bandit problem: after each new observation, the subject learns something about the bandit and thus transitions to a new belief state (MAB problems are thus more similar to Bayes-adaptive MDPs [2]). This instance of Need, therefore, quantifies how likely is the subject to ever encounter the potential update belief state according to its prior belief about each arm’s paoyff probability, as well as its current decision policy, *π*_old_.

We again assume a DYNA architecture whereby the subject may or may not decide to perform replay based on the expected improvement it estimates to its current decision policy. We set a threshold, *ξ*, which specifies the minimal EVB required for each potential replay update to be executed.

Fig S1B shows an example replay update which was executed first because the subject estimated it to provide the greatest improvement to the value of its root belief state. Moreover, this example also highlights the effect of generalisation due to each individual replay update: this is visible in how the Need term changes at all other belief states as a result of the single replay update. Fig S1C shows all replay updates executed by the subject for which the estimated benefit exceeded the fixed EVB threshold. Note that only 4 replay updates resulted in the accumulation of a near-optimal value at the subject’s root belief state, and that accumulated value reflected the benefit of future learning in an approximately optimal way (with respect to the subject’s prior belief). We additionally simulated our subject with varying parameters (such as the softmax inverse temperature, EVB threshold, and horizon), and those results are reported in Fig S2.

We further investigated the effect of behavioural policy on the statistics of the subject’s replay choices. This was done by randomly shuffling the true (fixed-horizon) initialised action values across all belief states before letting the subject engage in replay. This revealed that the initial state of knowledge of the subject (its behavioural policy) played a critical role in affecting the resulting benefit of replay (Fig S3) – which can furthermore be harmful [5]. This is visible from the wide distribution of the value of the resulting policy (Fig 3A), as well as the frequent lack of propagation of the value of distal beliefs towards the root of the tree (Fig 3B).

Finally, we examined the patterns of sequence and single-action replay updates (Fig S4). Our simulations indicated that the relative proportion of forward and reverse sequence replay was biased towards reverse replay (Fig S4); however, there was also a significant number of forward replay sequences (1-sample t test, *t* = 11.40, *p* ≪0.0001). Moreover, the total number of updated actions appeared to be greater with sequence replay compared to single-action replay updates (2-sample t test, *t* = 2.05, *p* = 0.042) which is expected given the open-loop nature of sequence replay optimisation. The full characterisation of sequence replay thus still remains an open question which we leave to future work.

### Implementation

#### Estimation of exploratory Need

We used a Monte-Carlo estimator for the Need term when calculating 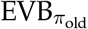 from equation 13 for determining the priority of replay updates. The subject’s belief space was discretised into its current belief state and two future possibilities for each of the uncertain barriers that they were either present or absent with certainty. Those possible belief states, moreover, could be envisioned by the subject only so long as they were within the reach of the subject’s limited horizon, *h*. We denote this limited horizon, discretised belief space as disc_*h*_(ℬ).

For the estimation of Need, *N* trajectories were simulated, all starting from the subject’s current belief state *b*_*ρ*_ = {*s*_*ρ*_, *P* (𝒫)}, where the decisions at each encountered belief state in each simulated trajectory were governed by the subject’s behavioural policy at those belief states and the belief state transitions – by the expected transition model associated with the subject’s belief state in the trajectory.

When attempting to cross one of the uncertain barriers in a given trajectory, the next belief state was sampled according to *b*^′^∼ 𝒯 (*b*^′^ |*b, a*). The subject’s belief about the transition dynamics in the new belief state, *b*^′^, was then updated according according to what actually happened. For successful transitions (with an open barrier), the probability of that transition was set to 1 with no remaining uncertainty; similarly, for failed transitions (with a closed barrier), the probability of that transition was set to 0, also with no remaining uncertainty.

All simulations were run so long as *γ*^*d*^, where *d* was the trajectory length, exceeded a fixed threshold, *ϵ* (which was always set to 10^−5^). Each *i*^th^ simulated trajectory returned the smallest number of steps, *K*_*i*_(*b*), that it took to reach each encountered belief state *b*∈ disc_*h*_(ℬ) along the trajectory, as well as the non-cumulative Need (time-discounted probability of reaching those belief states according to the belief transition model and the subject’s behavioural policy) upon the first encounter, 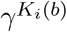, associated with those belief states.

Finally, for each encountered belief state *b* = {*s, P* (𝒫)} ∈disc_*h*_(ℬ), we estimated the Need using a second-form certainty equivalence. The subject accounted for the evolution of its prior belief up to the potential update belief state after which it assumed stationary transition model dynamics (and no forgetting). That is, the resulting Need was averaged over the non-cumulative Need encountered in each of *N* simulated trajectories, which accounted for the learning and transitions through belief states within the reach of the subject’s horizon, to which a certainty-equivalent Need was added with a stationary transition model of that belief state:

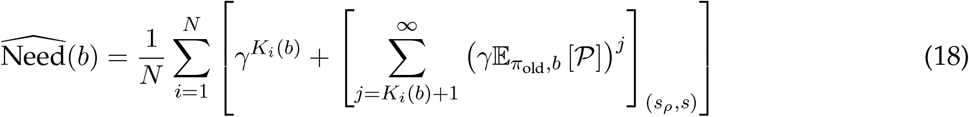

where [·]_(*i,j*)_ is a scalar value obtained by indexing the matrix by row *i* and column *j*.

Note that the expression 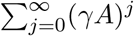, which corresponds to a geometric series for some matrix *A*, can also be written as (*I* −*γA*)^−1^. In Equation 18, however, the counter for the infinite matrix sum does not start at zero. This is because for the first *K*_*i*_(*b*) steps the transition model is non-stationary due to potential learning during those first steps within the reach of the subject’s horizon. After those first *K*_*i*_(*b*) steps, the subject computes the remaining of Need using the expected transition model of the final belief state in the simulated trajectory, 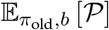.

#### Sequence generation

Sequence generation was implemented as an iterative procedure. All possible single-action updates were first generated, for belief states which were within the reach of the subject’s horizon – that is, all belief states in disc_*h*_(ℬ). Then, for forward sequences, all of the single-action updates were extended by applying all possible actions from the final belief state reached in those single-action updates (governed by the belief transition model 𝒯). This was repeated until sequences of the maximal specified length *L* were generated. Three important constraints we imposed on the sequence generation procedure: i) physical states encountered in the sequences were not allowed to repeat, hence preventing loops; ii) each sequence was extended by an additional action only if the 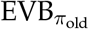 of the resulting sequence exceeded the 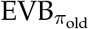 threshold; and iii) only those belief states contained in disc_*h*_(ℬ) were added to the sequences, such that the resulting sequences could not contain belief states outside of the subject’s horizon.

To generate reverse sequences, the same procedure was applied with the same imposed constraints. The only difference was the directionality of the value propagation along the action sequences. Note that the construction of reverse sequences requires an inverse belief transition model. An inverse transition model, for any given belief state *b*^′^, outputs a probability distribution over belief state-action pairs which quantifies how likely each of those are to result in a transition to *b*^′^. With our notation from Equation 12, given *b*^′^, an inverse transition model would assign zero probability to all belief state-action pairs but those for which *b*^′^ ∈*C*(*b, a*). When generating reverse sequences, we used a forward transition model (instead of learning a separate inverse transition model) which assigned the same uncertainty for reverse transitions as for forward ones.

#### Simulation details

Fig 1 was generated by simulating a vanilla Mattar & Daw [3] replay subject. The subject learned model-free *Q*-values according to equation 4, which it then used for online control through a softmax policy. We additionally imposed forgetting on the model-free *Q*-values learnt by the subject, after every move made by the subject, to imitate a continual learning problem such that replay remained throughout the whole simulated experiment [5]. The aforesaid forgetting was operationalised as the exponential decay towards the initialised values controlled by a forgetting parameter:

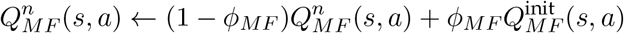

The state-transition model of the subject, *T*, was initialised such that it indicated that no barriers were present and the transition probabilities indicated the true transition structure. After every transition which attempted to cross the top-most barrier, the subject updated its state-transition model as:

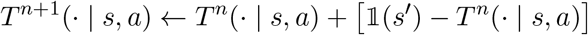

where 𝟙(*s*^′^) is a vector of the same dimension as the state space where each entry was zero except for the experienced next state, *s*^′^, for which the entry is 1. After every such update, the subject’s state-transition model probabilities associated with the uncertain barrier transition were normalised to ensure that they add up to 1:

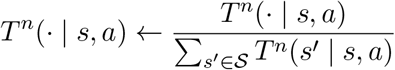

The subject additionally cached all observed experiences to use them in replay (we followed the same implementation protocol as in Mattar & Daw [3]). The memory buffer of the subject was updated after each corresponding online experience to account for the possible changes in the environment. The agent then engaged in replay after every move by prioritising the replay updates using equation 8 so long as the estimated 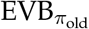 exceeded the minimal improvement threshold, *ξ*.

The subject was simulated for the first 2000 moves in the environment shown in Fig 1A-C. For the second 2000 moves, the environment was altered to that shown in Fig 1D-F without the subject being informed about such change. Note that the barrier was not bidirectional – the subject was not allowed to learn about the barrier from the state above it (i.e., it had to approach the barrier directly from the start state). The simulations were repeated 10 times and the average results are reported. The values of the free parameters used in those simulations are reported in Table 1.

**Table 1:**
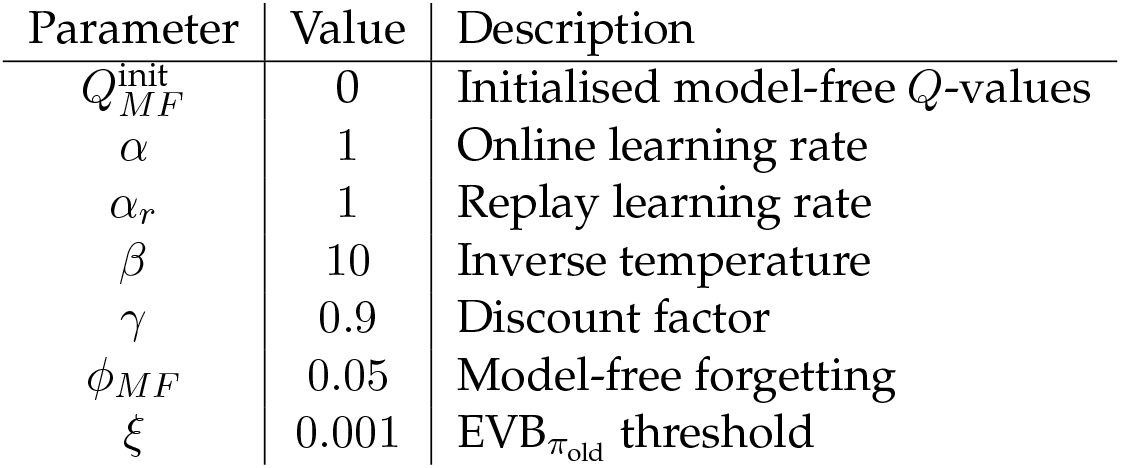
Simulation parameters for Fig 1.

Fig 2A was generated by performing regular value iteration with a transition model which assumed that both barriers were present. The tolerance threshold for value iteration was set to 10^−5^. The subject’s belief about the presence of the top barrier was then set to Beta(7, 2), after which it was allowed to engage in replay whilst being situated at the start state. The subject prioritised replay updates (equation 11) by calculating the Gain associated with all potential replay updates at all belief states within its horizon reach according to equation 13. The subject estimated Need for all potential replay updates using equation 18. Fig 2B-C show the prioritised replay updates and their order, as well as the new updated exploratory policy respectively.

For Fig 2D-E, the subject was situated at the state just below the top barrier. Its model-free *Q*-value for the action to cross the barrier was set to 0 to emulate the potential online discovery of the barrier being present; similarly, the subject’s belief was initialised to indicate the presence of the barrier with certainty. Accordingly, the subject’s belief was set to reflect the potential discovery of the barrier being present. The subject was then allowed to replay in the same way as described above. The values of the free parameters used in the shown simulations are reported in Table 2. In this and all subsequent tables reporting the parameter values, we highlighted the crucial parameters and their values which differed between the simulations.

**Table 2:** Simulation parameters for Fig 2.

| Parameter | Value | Description |
| --- | --- | --- |
| $\alpha$ | 1 | Online learning rate |
| $\alpha_r$ | 1 | Replay learning rate |
| $\beta$ | 2 | Inverse temperature |
| $\gamma$ | 0.9 | Discount factor |
| $\alpha_T, \beta_T$ | 7, 2 | Beta prior parameters for the top barrier |
| $h$ | 8 | Planning (replay) horizon |
| $N$ | 2000 | Number of simulated trajectories |
| $\xi$ | 0.001 | $\text{EVB}_{\pi_{\text{old}}}$ threshold |

Fig 3 was generated in the same way as Fig 2 but the replays that the agent was allowed to execute additionally included sequence events. The maximal sequence length, *L*, was constrained to be the distance between the start state and the uncertain barrier. The agent prioritised which replay updates to execute by choosing from all possible replay updates of lengths 1 through *L*. The online discovery was operationalised in the same way as in Fig 2, and the replay process was then repeated with the subject being situated in the new belief state. The values of the free parameters used in the shown simulations are reported in Table 3.

**Table 3:**
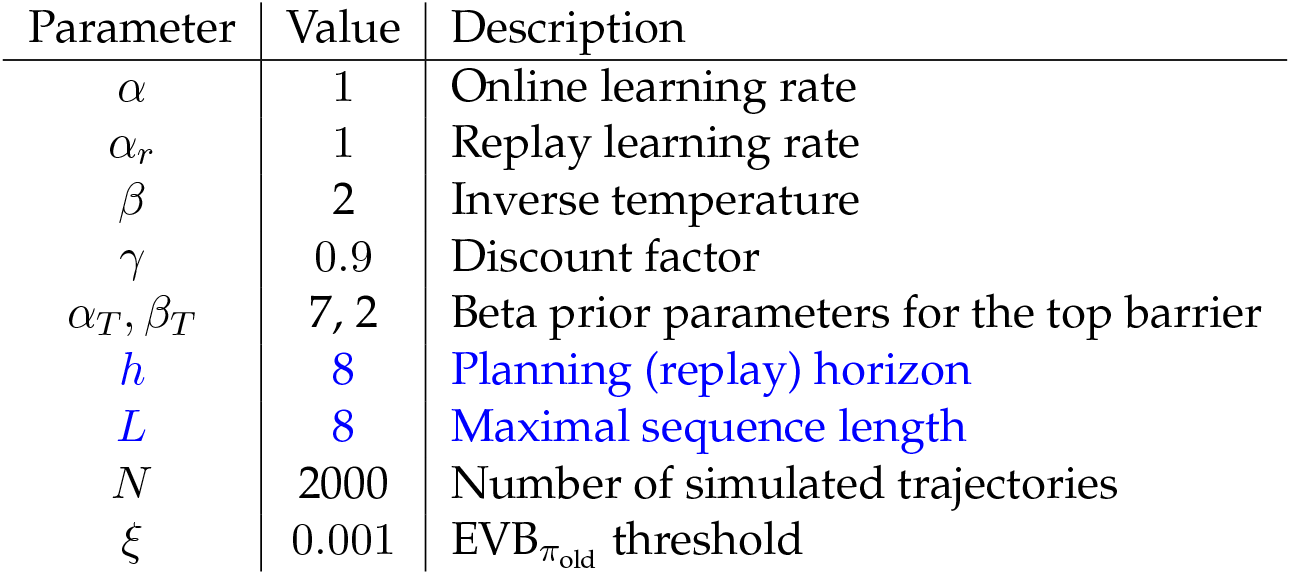
Simulation parameters for Fig 3.

Fig 4A-C was generated in the same way as Fig 3 except that the subject was uncertain about two barriers at the same time. The parameter values used in the shown simulations are reported in Table 4. For Fig 4D, the *Q*-values were initialised to 0. First, the subject was allowed to replay with the knowledge that reward was absent in both arms. Next, it was allowed to replay with the knowledge that the right (‘cued’) arm contained reward. All other simulation parameters were kept the same except the planning horizon which was set to 3.

**Table 4:** Simulation parameters for Fig 4.

| Parameter | Value | Description |
| --- | --- | --- |
| $\alpha$ | 1 | Online learning rate |
| $\alpha_r$ | 1 | Replay learning rate |
| $\beta$ | 2 | Inverse temperature |
| $\gamma$ | 0.9 | Discount factor |
| $\alpha_T, \beta_T$ | 7, 2 | Beta prior parameters for the top barrier |
| $\alpha_B, \beta_B$ | 7, 2 | Beta prior parameters for the bottom barrier |
| $h$ | 6 | Planning (replay) horizon |
| $L$ | 4 | Maximal sequence length |
| $N$ | 2000 | Number of simulated trajectories |
| $\xi$ | 0.001 | $\text{EVB}_{\pi_{\text{old}}}$ threshold |

**Table 5:** Simulation parameters for Fig S1.

| Parameter | Value | Description |
| --- | --- | --- |
| $\alpha_r$ | 1 | Replay learning rate |
| $\beta$ | 4 | Inverse temperature |
| $\gamma$ | 0.9 | Discount factor |
| $\alpha_1, \beta_1$ | 5, 3 | Beta prior parameters for arm 1 |
| $\alpha_2, \beta_2$ | 1, 5 | Beta prior parameters for arm 2 |
| $h$ | 2 | Planning (replay) horizon |
| $\xi$ | 0.01 | $\text{EVB}_{\pi_{\text{old}}}$ threshold |

Fig S1 was generated by constructing a belief tree of horizon 2 which was rooted at the subject’s prior belief about the payoff probabilities of the two arms. The *Q*-values for all actions in all belief states were initialised to 0, except for those at the final horizon which were initialised to the expected immediate payoff according to those beliefs. Gain associated with each replay update in the tree (equation 11) was calculated according to equation 17 and Need associated with every update belief state was calculated according to equation 17.

Fig S2 was generated by varying the subject’s policy, the 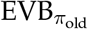 threshold, as well as the subject’s planning horizon (all values are reported in the figure). The root value shown was taken as the expected return the subject expected at the root belief after all executed replay updates. The value of the evaluated policy was computed by evaluating the new updated policy as a result of the replay updates in the whole tree.

Fig S3 was generated in the same way as Fig S2 but the results were averaged over 200 random value initialisations in the tree. The randomisation was achieved by first performing full value iteration in the tree, and hence computing the true fixed-horizon values associated with each action in the tree. Next, those values were randomly shuffled across all belief states in the tree. Fig S3 shows the average, as well as individual replay processes in the randomised trees.

The data shown in Fig S4 was generated in the same way as for Fig S3, as well as additionally allowing the subject to perform sequence replay where the maximal sequence length was constrained to the horizon of the tree.

Fig S5 were generated in the same way as Fig 3 but the subject was initialised with a different prior belief about the presence of the barrier. In this case, the prior belief was set to Beta(2, 2).

The data in Fig S6 were generated in the same way as for Fig 3 but the subject was initialised with a range of different prior beliefs about the presence of the barrier.

Fig S7 was generated with the same parameter values as Fig 4 (shown in Table 4) but with the planning horizon set to 12.

## 3 Supplementary figures

**Figure S1:**
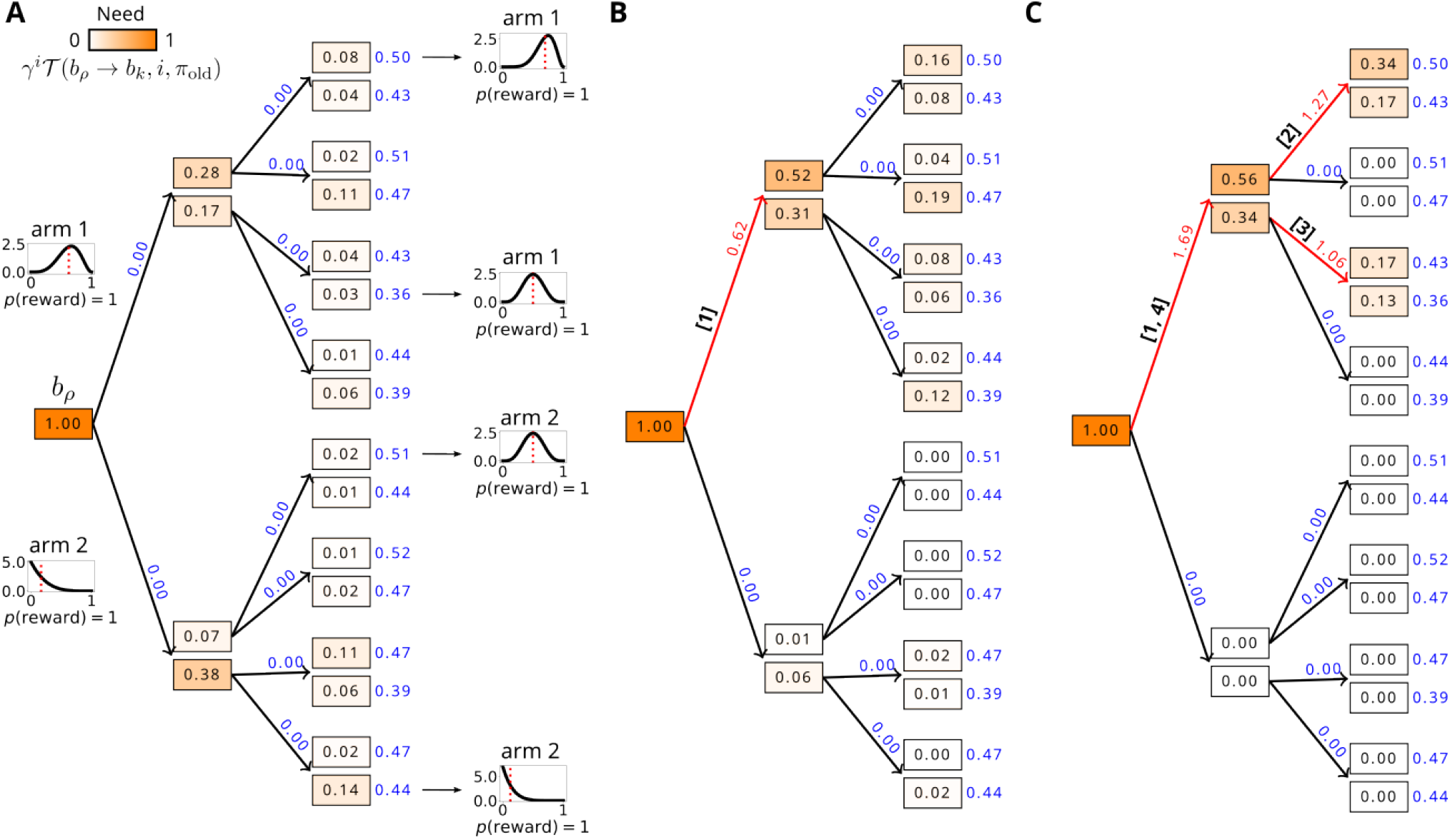
Replay updates in Bandit belief space. A) Planning tree of horizon 2. Each rectangle corresponds to a distinct belief state. The leftmost belief state, at the root of the tree, corresponds to the subject’s prior belief, *b*_*ρ*_. The insets next to some belief states graphically demonstrate the subject’s belief about the payoff of one of the arms in those belief states (the red dotted lines show the resulting mean payoffs). For the paired belief states, the top ones always result from imagined successful outcomes (received a reward of 1), whereas the bottoms ones – from imagined failed outcomes (received a reward of 0). Belief states are coloured according to their exploratory Need; moreover, Need is additionally shown with numbers in each belief state. Since the subject’s behavioural policy is stochastic (softmax), all belief states have positive estimated Need (for some belief states, it is shown as 0.00 for demonstration purposes since those values were too small). The black arrows show actions available at each belief state. The top arrows always denote the choice of arm 1 and the bottom arrows – arm 2. The blue numbers above each action arrow denote the *Q*-values associated with each action in every belief state. All *Q*-values were initialised to 0 except for those belief states at the final horizon for which the initialisation values were determined by the expected immediate reward according to those belief states. B) Single replay update in the belief tree. The subject chose to update the *Q*-value of arm 1 at the prior belief state (the updated action arrow is highlighted in red) towards the expected value of the two belief states at the next horizon (the new updated value is highlighted in red). This replay update was executed because i) it was estimated to have the greatest EVB; and ii) the estimated EVB of this update exceeded the EVB threshold. Note the effect of generalisation of this individual replay update which is visible in how the Need that the subject calculates for all other belief states changes throughout the tree. C) All replay updates executed by the subject until the estimated benefit was calculated to be below the EVB threshold. The bold numbers in squared brackets show the order in which those updates were executed. The action values highlighted in red are the final action values updated by all shown replay updates in the tree.

**Figure S2:**
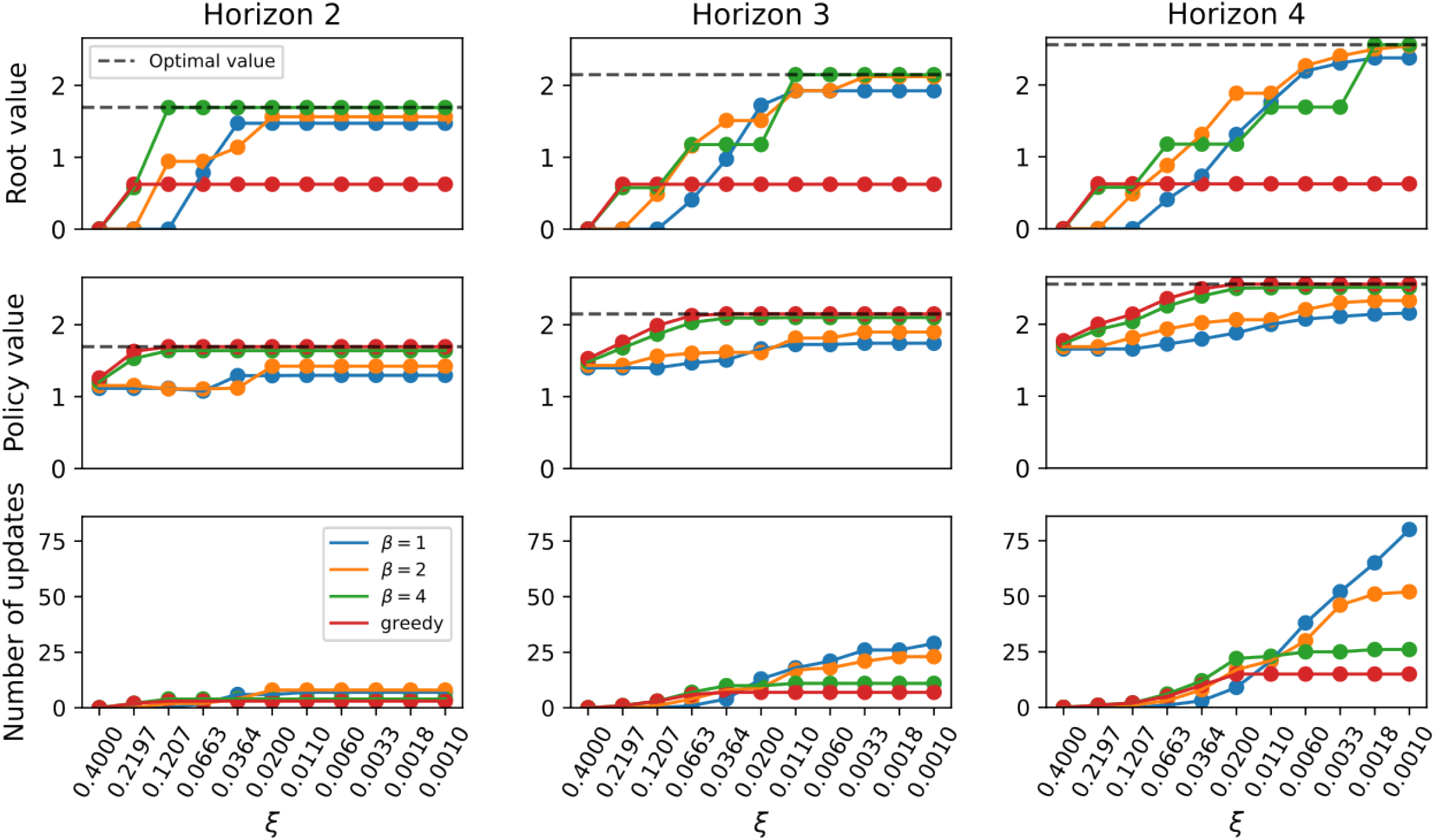
Policy improvement occasioned by replay. Top: Evolution of the value of the root belief state in the bandit task (same as in Fig S1) due to replay as a function of the EVB threhsold, *ξ*. Middle: Evolution of the value of the policy (evaluated in the belief tree) which resulted from replay updates at different EVB thresholds. Bottom: Total number of replay updates executed for the different EVB thresholds.

**Figure S3:**
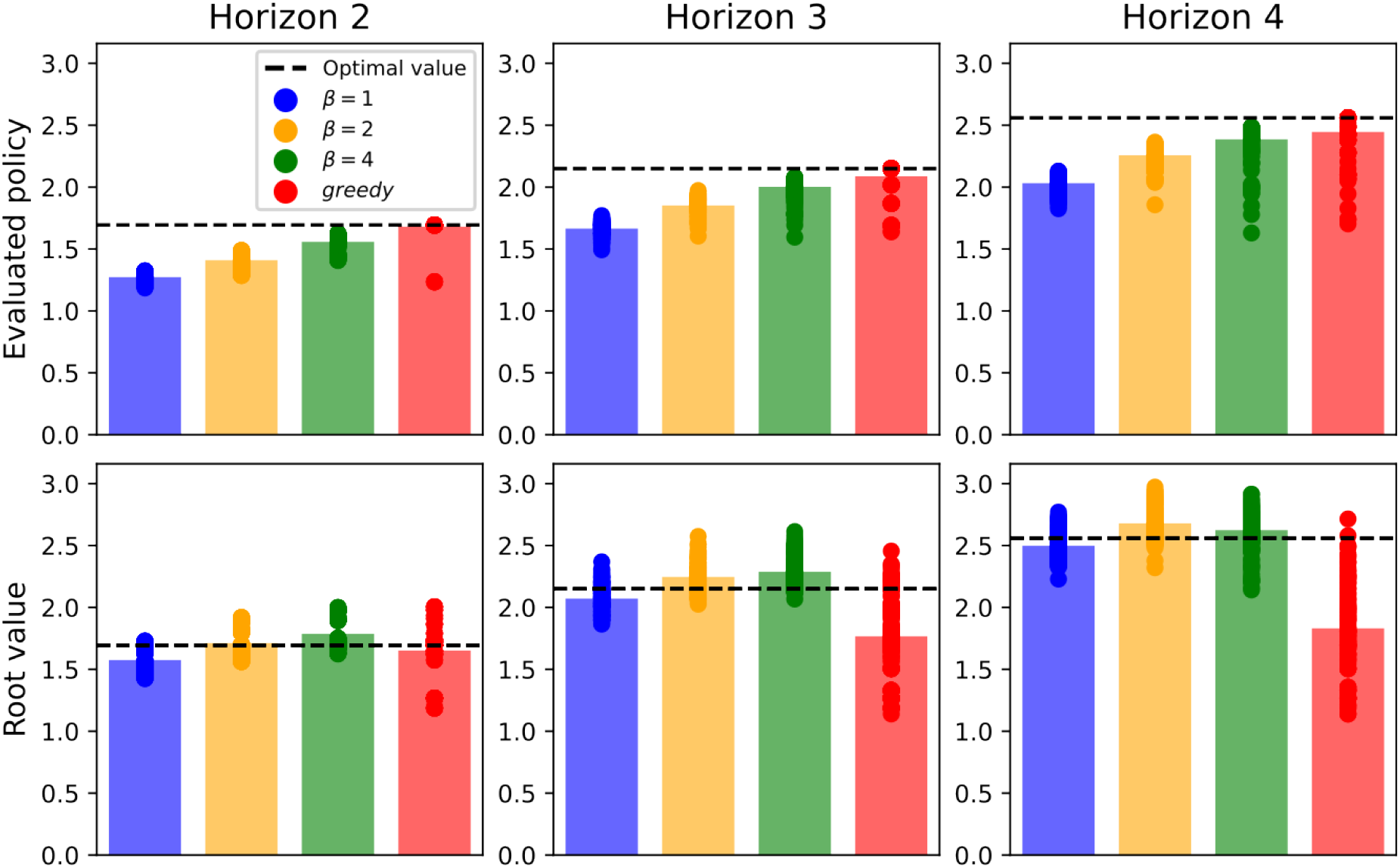
Effect of initialised behavioural policy. Top: The final value of the root belief state in the bandit task (same as in Fig S1) due to replay with a fixed EVB threshold. The initialised values of all belief states were randomised to imitate noisy initial experience (or potential changes in the bandit payoff probabilities). The bars show average root belief state values over 200 different tree initialisations. Each dot corresponds to an individual tree. Bottom: Same as above but for the value of the updated policy evaluated in the tree.

**Figure S4:**
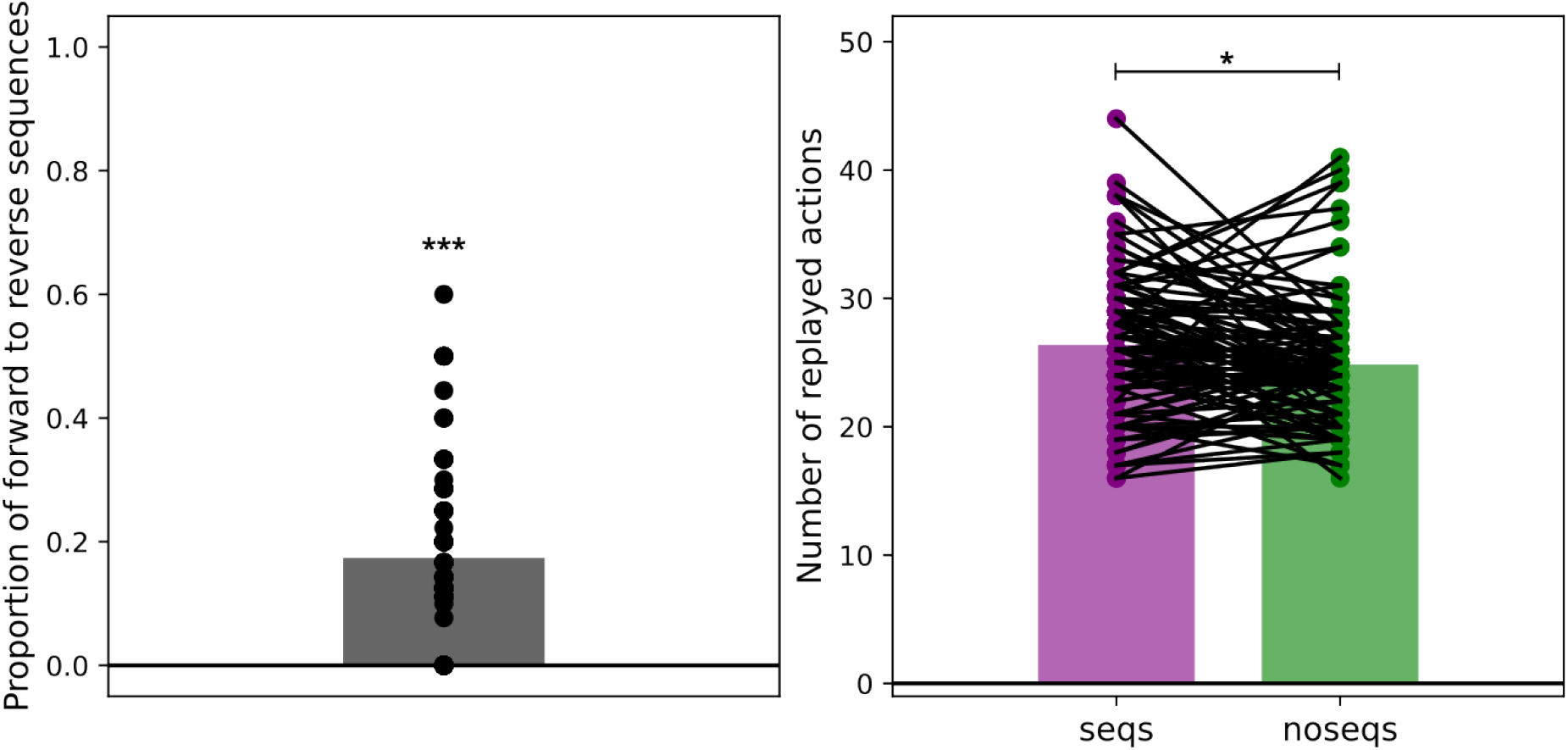
Sequence replay statistics. Left: Proportion of forward to reverse sequences replayed in the belief tree in the bandit task with the same prior belief as in Fig S1 with planning horizon set to 4. The initialised values of all belief states were randomised as in Fig S3. The bar shows average proportion over 200 different tree initialisations. Right: Average number of replayed actions in the same tree initialisations as above with and without sequence replay. *** *p <* 0.001, * *p <* 0.05.

**Figure S5:**
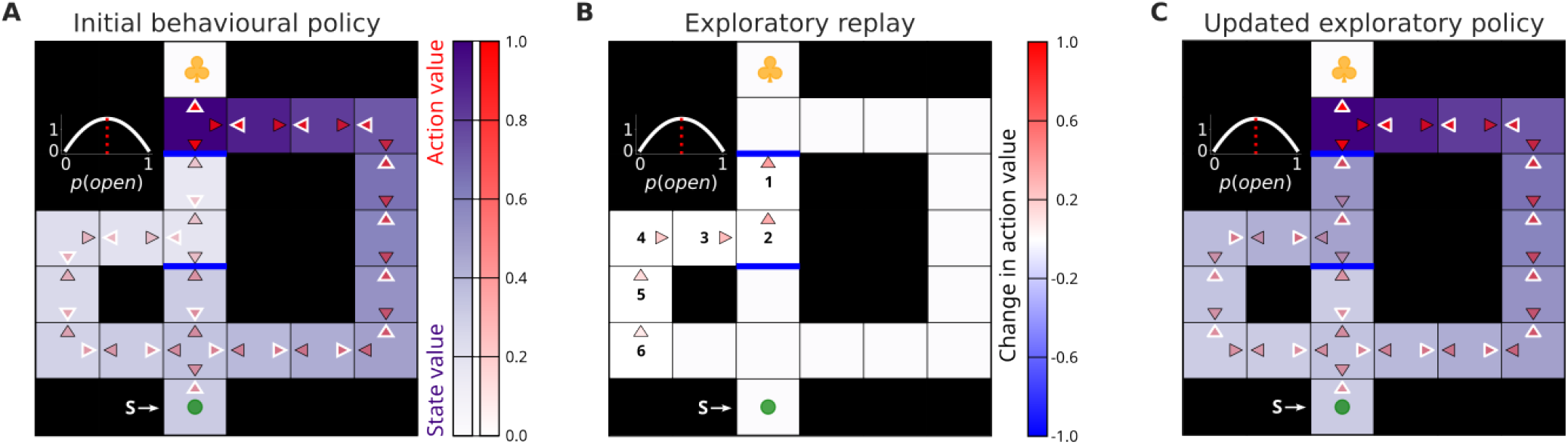
Uncertainty affects replay choices and their behavioural readout. The layout of the figure is similar to that of Fig 2. A) Prior state of knowledge of the subject. In this example, the subject’s belief was more pessimistic since it indicated a lower subjective probability of the top barrier being potentially open (as evident from the expected probability the subject accorded to this possibility, shown with the red dotted line in the inset). B) The value of exploration was estimated to be lower (since the subject’s belief was more pessimistic), and therefore replay did not propagate the benefit of exploration deep enough (towards the subject’s location). This is in part owing to the temporal discounting which decays the benefit of exploration with travel distance. C) The updated policy still prescribed the subject to exploit the longer path (maximal *Q*-values at each state are again shown with white outlines), since the critical action at the junction between the different arms had not been updated by exploratory replay.

**Figure S6:**
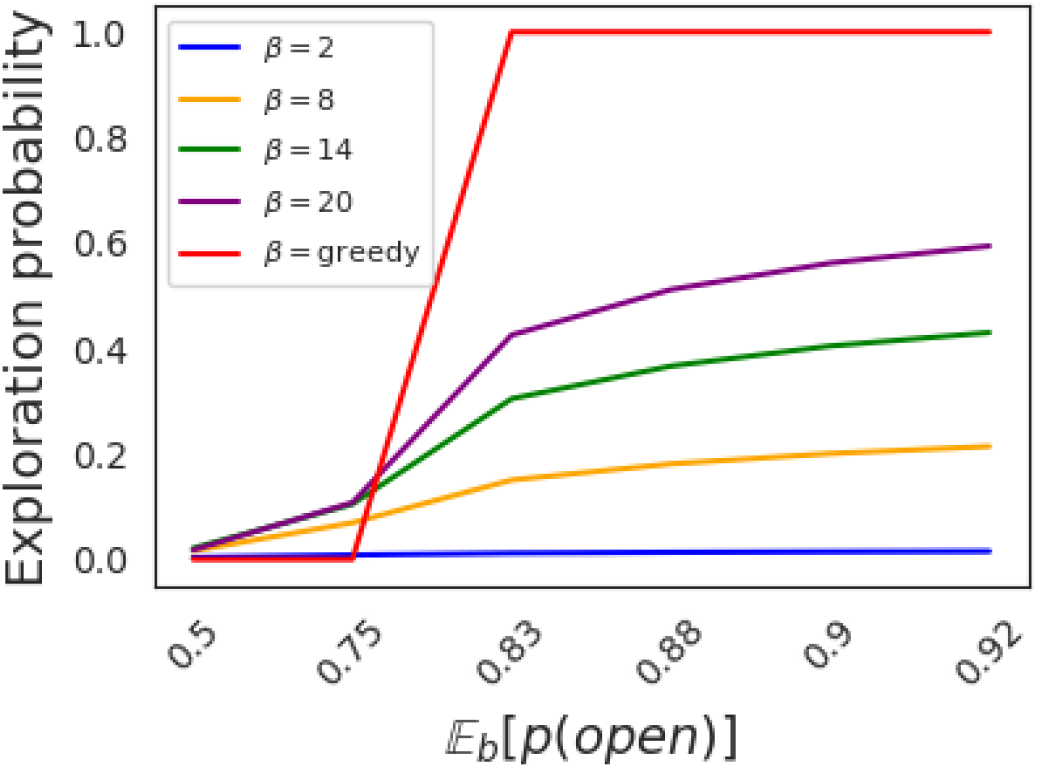
Relationship between uncertainty, behavioural policy and exploration quality. The graph shows the marginal probability of directed exploration (approaching and attempting the potential barrier in Figs 2 and 3 from the start state) as a function of the subject’s uncertainty and the greediness of its behavioural policy. As the subject’s belief (𝔼 [*p*(open)]) in the absence of the barrier increased, it became progressively more likely to engage in the act of directed exploration. The same softmax policy with inverse temperature *β* = 2 was used to calculate the priority of replay updates. However, applying different inverse temperature parameters (which subjects might heuristically use to arrange for offline exploration) to the resulting exploratory value function yielded policies with different incentives for exploration.

**Figure S7:**
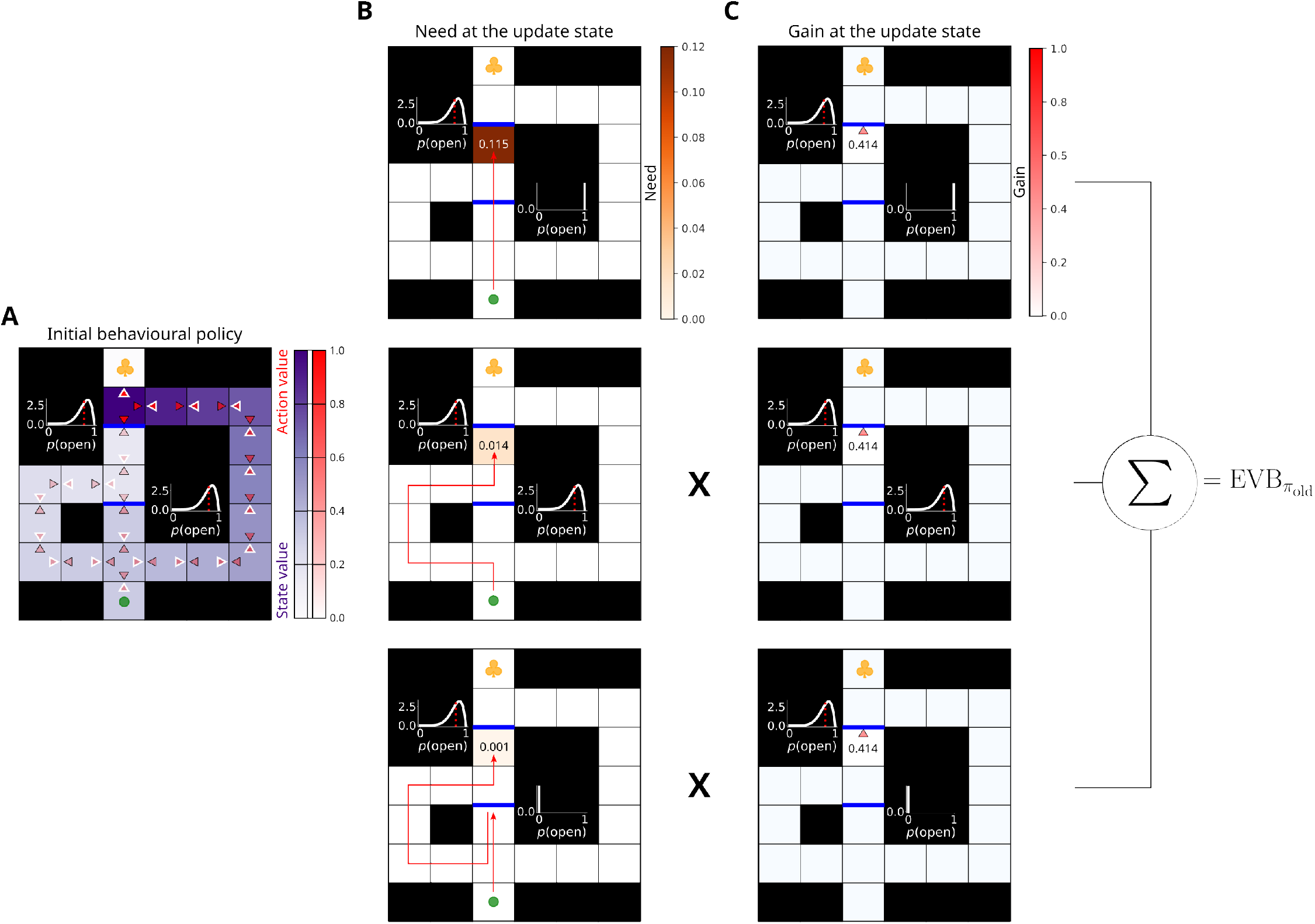
The benefit of generalisation in replay across belief states. A) Prior state of knowledge of the subject. The layout of the panel is identical to that of Fig 2. B) Need that the subject estimated for the potential update at the physical state below the top-most barrier. Each row shows a different belief with which the subject can reach that physical state. The red arrows denote the potential routes to that physical location that the agent can undertake all of which result in different belief states. For brevity, we only show a restricted number of the possible (discretised) beliefs. C) Estimated gain for the potential update of the action that attempts to cross the barrier. Note that Gain is positive in all the shown belief states associated with the top-most barrier. This means that the subject can expect to accrue more reward due to the update at that physical location whilst reaching it with different beliefs about the other (bottom) barrier. This knowledge of the potential future beliefs allows the subject to generalise across belief information states.

**Figure S8:**
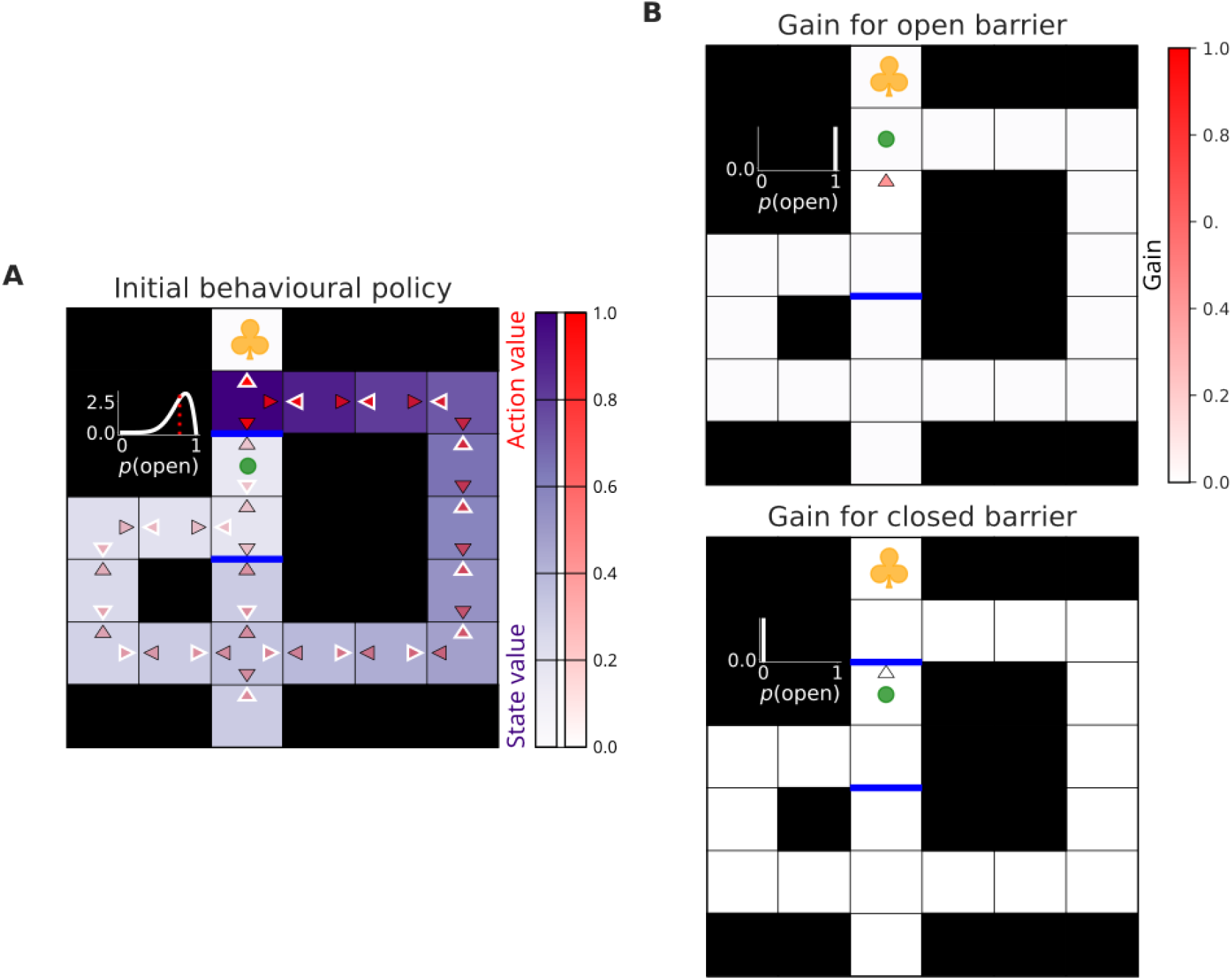
Exploratory Gain. A) Prior state of knowledge of the subject. The layout of the panel is identical to that of Fig 2A. B) Top: Gain that the subject estimates as a result of the imagined successful shortcut transition through the potential barrier just above the subject. Bottom: similar to the above but Gain estimated for the imagined failed transition through the potential barrier. The full exploratory Gain is then calculated as the expected Gain of the two possible outcomes above weighted by their respective prior probabilities determined by the subject’s belief state in A).

## 4 Acknowledgements

The authors thank Philipp Schwartenbeck, David Foster, Christopher Gagne, Mihály Bányai, and Noa Hedrich for their valuable feedback on the manuscript. Philipp Schwartenbeck and Christopher Gagne additionally contributed to the earlier ideas relevant to this work.

